# Neural Signatures of Post-Decision Outcome Expectation and Evaluation in Human Sensorimotor Choice Behavior

**DOI:** 10.64898/2026.01.23.701339

**Authors:** Niloofar Gharesi, John F Kalaska, Sylvain Baillet

**Affiliations:** McConnell Brain Imaging Centre, Montréal Neurological Institute, McGill University, Montréal, Canada; Groupe de recherche sur la signalisation neuronale et la circuiterie, Département de neurosciences, Université de Montréal, Montréal, QC, Canada; Centre de recherche du Centre hospitalier de l’Université de Montréal (CRCHUM), Montréal, QC, Canada; Département de neurosciences, Université de Montréal, Montréal, QC, Canada

## Abstract

The concept of embodied sensorimotor decision-making proposes that processes implicated in evaluating sensory inputs and selecting appropriate motor actions unfold partly in cortical regions traditionally associated with movement planning and execution. Reinforcement learning models emphasize the role of reward prediction error (RPE) in optimizing action selection based on decision outcome feedback. However, most evidence for the existence of RPE signals locates them in midline frontal and parietal cortex, and comes from tasks with externally manipulated reward probabilities that create artificial prediction errors. Whether RPE signals are expressed in human cortical motor areas during deterministic (non-probabilistic) tasks remains unclear, and would provide further support for embodied decision-making.

We used magnetoencephalography (MEG) to study post-decision neural dynamics in a color discrimination task in selected cortical regions of interest (ROIs). Participants had to press buttons with their left or right index finger in response to checkerboard stimuli with different levels of color evidence for the correct choice. Outcomes were fully determined by participants’ choices. Delayed auditory feedback veridically indicated whether their hand choice was correct or not.

We observed a robust beta-band (15–29 Hz) rebound after correct outcome feedback, strongest in ventral and dorsal premotor, anterior cingulate and superior parietal ROIs as well as occipital and auditory ROIs, and weakest in the primary motor and somatosensory ROIs. Critically, the rebound magnitude after correct feedback scaled inversely with color evidence strength and associated decision error rates. It was minimal in strong-evidence trials (∼0.1% errors) and maximal in weak-evidence trials (∼34% errors), resembling a context-sensitive positive RPE signal that was strongest when a correct outcome was least expected. Alpha-band (8–12 Hz) post-feedback rebound increases in weak evidence trials were not as strong as in the beta band and appeared mainly in occipital, superior parietal and posterior cingulate ROIs.

After the decision but before feedback, both beta and alpha band power showed sensitivity to the level of sensory evidence on which the decisions had been based, with reduced post-movement rebound or enhanced suppression in trials with weak evidence—suggestive of internally generated outcome expectations. Pre-feedback alpha rebound suppression was strongest in occipital, superior parietal and posterior cingulate ROIs. Pre-feedback beta rebound suppression was not as strong.

Together, these findings reveal distinct beta- and alpha-band dynamics that reflect internal pre-feedback outcome expectations and feedback-driven RPE-like outcome assessments. They support distributed cortical mechanisms, including premotor, parietal and cingulate regions, in reward expectation, outcome evaluation, and adaptive control, highlighting a role for motor and associative cortices in embodied decision-making, performance monitoring, and flexible behavior under uncertainty.

## Introduction

Perceptual (or sensorimotor) decision-making has been extensively studied in non-human primates (NHPs) in tasks that required the NHPs to evaluate sensory inputs and execute motor responses according to sensory-motor association rules (Brody and Hanks 2016; Cisek and Kalaska 2010; Gold and Shadlen 2007; Hanks and Summerfield 2017; Kable and Glimcher 2009; Roitman and Shadlen 2002; Romo and de Lafuente 2013; Shadlen and Kiani 2013).

A series of studies particularly relevant to the present work required participants to select one of two color-coded targets based on either a monochromatic cue (Cisek and Kalaska 2005; Coallier and Kalaska 2014) or a perceptual decision about the dominant color of a multi-colored checkerboard-like (CKB) stimulus (Boucher et al. 2023; Chandrasekaran et al. 2017; 2019; Coallier et al. 2015; Coallier and Kalaska 2014; Genkin et al. 2023; Löffler et al. 2023; Mante et al. 2013; Wang et al. 2019). The CKB paradigms varied decision difficulty and decision outcome accuracy by changing the relative number of squares of two colors in the CKB, forming the conceptual basis for the current study.

Behavioral and neural results from such tasks strongly support “integrate-to-threshold” decision models, in which sensory evidence accumulates until a decision threshold is crossed, triggering final motor action selection and execution (Cisek and Kalaska 2010; Ditterich 2006; Gold and Shadlen 2007; Mazurek et al. 2003; Shadlen and Kiani 2013). While classical cognitive theories posit that decisions occur in an abstract “cognitive space” before activating the motor system (Fodor 1983; Pylyshyn 1984), the single-neuron recordings challenged this distinction. Instead, they showed that decision-related variables, including evidence strength, ambiguity, and available action alternatives, are expressed in motor-related cortical regions before the final action choice is made and executed (Churchland et al. 2008; Cisek and Kalaska 2005; Klaes et al. 2011; Platt and Glimcher 1999; Roitman and Shadlen 2002; Thura et al. 2022; Thura and Cisek 2014; Augusto et al. 2025). These results support the embodied decision-making framework, in which sensorimotor decision processes unfold partially within circuits traditionally implicated in action execution (Cisek and Kalaska 2010; Cisek and Pastor-Bernier 2014; Gharesi et al. 2023; Gold and Shadlen 2007; Gordon et al. 2021).

Additional evidence for embodiment comes from studies showing that motor cortical activity prior to or during movement is modulated by potential gains and reward expectation (An et al. 2019; Marsh et al. 2015; Roesch and Olson 2003). Cortical motor areas also respond differentially to unexpected reward delivery or omission after a movement, suggesting a role in performance monitoring and action outcome evaluation (Ramakrishnan et al. 2017; Ramkumar et al. 2016).

A central concept in outcome monitoring is the Reward Prediction Error (RPE), defined in Reinforcement Learning (RL) theory as the discrepancy between the expected and experienced outcome of a decision or an action (Barto and Sutton 1982; Holroyd and Coles 2002; Sutton and Barto 1981; Montague et al. 1996). RL theory distinguishes two properties of the scalar value of an RPE. Its signed valence (positive or negative) indicates that the experienced outcome was better or worse than expected, whereas its unsigned magnitude indicates the degree of salience or “surprise” of the discrepancy, from zero (fully expected outcome) to large (a very unexpected outcome) (Holroyd and Coles, 2002; Ferdinand et al., 2012; Padron et al., 2016; Fouragnan et al., 2018; Walentowska et al., 2019; Hoy et al., 2021, 2023). A positive RPE indicates that the experienced outcome exceeded expectation, promoting reinforcement of the preceding action, whereas a negative RPE suggests the opposite, discouraging similar future behavior. The unsigned magnitude could influence the rate at which those adaptations are applied, as well as the level of arousal or attention that might be evoked by the experienced outcome feedback.

The principal objective of this study was to provide further support for the embodied decision-making hypothesis by identifying magnetoencephalography (MEG) correlates of RPE signals in human cortical sensorimotor areas evoked by outcome feedback cues in deterministic decision-making tasks with ambiguous colored CKB stimuli that presented different amounts of sensory evidence for two motor responses (Boucher et al. 2023; Chandrasekaran et al. 2017; 2019; Coallier et al. 2015; Coallier and Kalaska 2014; Genkin et al. 2023; Löffler et al. 2023; Mante et al. 2013; Wang et al. 2019). However, as in many other studies, we did not test all the formal criteria for an RPE signal as defined in RL theory (e.g., Montague et al., 1996; Schultz et al., 1997; Fiorillo et al., 2003; Talmi et al., 2012), and we present here only the responses to feedback cues that signalled correct decisions. Therefore, we will use the more phenomenological terms “RPE-like signal” to refer to any neural response to an outcome feedback cue that was modulated by outcome expectation, and “positive RPE-like signal” (“+RPE-like”) to refer specifically to an outcome-feedback-locked neural response whose magnitude scaled inversely with the expectation of a correct outcome, rather than to a fully signed RPE as defined in formal RL models.

The existence of RPE computations in the brain is well supported by neurophysiological studies. Midbrain dopaminergic (DA) neurons express RPE signals (Montague et al. 1996; Schultz et al. 1997). When reward probability is directly manipulated experimentally, they respond strongly to an unexpected reward when the instructed probability is low, but weakly when reward probability is high and a reward is expected (Schultz et al. 1997; Fiorillo et al. 2003). In sensorimotor decision-making tasks, DA responses to rewards received after correct decisions are stronger in trials in which sensory evidence supporting the correct choice was weak than in trials in which evidence was strong, and DA activity before reward delivery reflected critical information needed for RPE computations, including sensory evidence, perceptual uncertainty, decision confidence and reward expectation (Nomoto et al. 2010; Lak et al. 2017; de Lafuente and Romo 2011; Sarno et al. 2017).

Only a few single-neuron studies have searched for RPE-like signals in cortical motor regions. Wise and colleagues (Brasted and Wise 2004; Buch et al. 2006) showed that dorsal premotor cortex (PMd) neurons in NHPs express +RPE-like responses during visuomotor learning of new stimulus-response mappings between 4 novel visual icons and 4 hand movements. Initially, when the NHPs made the correct movement in response to an icon by chance, many PMd neurons responded strongly to reward delivery. As performance improved, reward-related PMd responses decreased progressively, consistent with a diminishing RPE-like signal as correct outcomes became increasingly expected. Neurons in the Supplementary Eye Field (SEF) express signals reflecting decision confidence before outcome feedback (So and Stuphorn 2016) and RPE-like responses after outcome feedback (So and Stuphorn 2012) in an oculomotor gambling task in which monkeys chose between two colored targets that offered different reward sizes and probabilities of reward delivery. Ramakrishnan et al. (2017) reported RPE-like responses to rewards at the end of simple instructed-reach movements. However, reward expectation and RPE were manipulated in that study by varying the timing of the reward after each reach, and by deliberately omitting the reward or delivering a double-sized reward in random trials, rather than by manipulating evidence to alter choice accuracy. Moreover, the sign of their RPE-like responses was inverted, being smaller with unexpected double-sized rewards. Finally, neural correlates of action outcomes (Levy et al. 2020) and RPE-like signals (Makino and Suhaimi 2025) are expressed in rodent sensorimotor cortex during skilled motor learning tasks.

In humans, EEG studies have revealed cortical signatures of RPE-like computations in the movement error-related negativity (ERN) and in outcome feedback-evoked potentials including the feedback-related negativity (FRN) and the later P3/P300 potential. Holroyd and Coles (2002) demonstrated that the ERN scales with expectancy of correct or incorrect outcomes, supporting its role in RPE computation. In a gambling task, Holroyd et al. (2003) showed that the FRN and associated P3/P300 are stronger when an expected reward was unexpectedly omitted probabilistically. Evidence for both positive and negative RPE-like responses have been reported across multiple task paradigms that used a range of different outcome feedback policies (Gehring et al. 1993; Miltner et al. 1997; Holroyd et al. 2003; 2004; Hajcak et al. 2005; Holroyd and Krigolson 2007; Hajcak et al. 2007; Holroyd et al. 2008; Yasuda et al. 2004; Frank et al. 2005; Cohen et al. 2009; 2007; Cohen and Ranganath 2007; Marco-Pallarés et al. 2008; Philiastides et al. 2010; Haji-Hosseini et al. 2012; HajiHosseini and Holroyd 2015a, b; Hoy et al. 2021, 2023; Billeke et al. 2020; Walentowska et al. 2019; Yaple et al. 2018; Fouragnan et al. 2017; Weismüller and Bellebaum 2016; Bai et al. 2015; Ferdinand et al. 2012; San Martín et al. 2010).

A smaller number of MEG studies have also reported RPE-like signals (Miltner et al. 2003; Steffen et al. 2011; Bunzeck et al. 2011; Apitz and Bunzeck 2012, 2014; Thomas et al. 2013; Bach et al. 2017). Doñamayor et al. (2011) identified an MEG FRN analog at 230–465ms post-feedback, sensitive to outcome valence (correct vs incorrect) and loss magnitude in parietal and caudal anterior cingulate cortex, with stronger beta-band responses (18–40 Hz) during high-reward trials. Doñamayor et al. (2012) found anticipatory and post-feedback beta/theta oscillations linked to reward magnitude and expectancy. Talmi et al. (2012) observed a fronto-central signal scaling with outcome valence and expectation in compliance with formal axiomatic criteria for an RPE in a probabilistic gambling task.

These EEG and MEG results showed that reward expectation strongly modulated neural responses to trial outcome feedback, demonstrating that cortical systems express temporally and spectrally specific RPE-like signals across frontoparietal and midline cortex to support adaptive behavior. However, prior human MEG and EEG studies have rarely examined RPE correlates in cortical motor areas (Cohen and Ranganath 2007). Moreover, there is little consensus in the field about the details of these RPE-like signals, including their sensitivity to outcome valence (either only positive or negative outcomes, or both), and their sensitivity to the magnitude of the difference between expected and experienced outcomes (either only positive or negative differences, or both) (e.g., HajiHosseini and Holroyd 2015a, b; Hoy et al. 2021, 2023). Indeed, a hallmark of this literature is that the diversity of findings is as large as the diversity of task designs and data analyses that were used.

Furthermore, many of the EEG and MEG studies that have reported RPE-like effects have relied on speeded-response tasks to induce performance errors, or on gambling, “guessing” or other response-choice tasks that used experimentally manipulated probabilistic reward policies in which the degree to which an outcome is expected or unexpected is determined in part or entirely by the level of capriciousness of the policy that determines the valence (e.g., correct/incorrect, or gain/loss) and the magnitude (large or small gains or losses) of the outcome of each trial, independent of the participant’s own decisions. These paradigms may not generalize to many real-world sensorimotor decisions, where outcomes depend only on the accuracy of the participant’s internal evaluation of sensory evidence. If RPEs drive naturalistic learning of sensorimotor decisions leading to action selection, they should appear in tasks where outcomes are determined solely by participants’ decisions, as has been reported in some prior studies (Miltner et al. 1997; Holroyd and Krigolson 2007; Holroyd et al. 2008; Ferdinand et al. 2012; Pardo-Vazquez et al. 2008; 2014; Padrón et al. 2016; Hoy et al. 2021, 2023).

We used MEG to examine post-decision neural activity in a priori defined sensorimotor cortical regions of interest during a CKB color discrimination task (Coallier and Kalaska 2014; Coallier et al. 2015; Wang et al. 2019) involving bilateral hand responses to report the perceptual decisions. Critically, outcomes were determined solely by the participants’ decisions, and delayed auditory feedback delivered after each decision informed them accurately of a correct or incorrect outcome. This enabled analysis of post-feedback responses in correct trials as potential +RPE-like signals based solely on the participant’s own introspective metacognitive evaluation of task performance and outcome expectations. We focused here on changes in beta and alpha band power during the post-decision epochs of the tasks.

A pre-movement suppression of beta band power followed by a post-movement beta rebound in sensorimotor cortex has been well documented in many EEG, MEG and LFP studies and has been attributed multiple roles in motor planning and execution (Rogge et al. 2022; Jurkiewicz et al. 2006; Pfurtscheller and Lopes Da Silva 1999; Engel and Fries 2010; Cheyne 2013; Barone and Rossiter 2021; An et al. 2019; Chouinard and Paus 2006; Kilavik et al. 2013; Tan et al. 2016). Changes in beta-band power have also been documented during several different processing stages in sensorimotor decision-making tasks, including correlations with sensory evidence, categorical perceptual decisions and motor-report choices (Donner et al. 2007; 2009; Cheyne et al. 2012; Lange et al. 2013; O’Connell et al. 2012; Kelly and O’Connell 2013; Twomey et al. 2016; Haegens et al. 2010; Spitzer et al. 2010; 2014; Herding et al. 2016; Rogge et al. 2022; Chandrasekaran et al. 2017; 2019; Haegens et al. 2011a).

Alpha/mu band activity is prominent in parieto-occipital and sensorimotor regions during sensorimotor decision-making tasks (Haegens et al. 2010; Haegens et al. 2011b). It often displays a movement-related suppression/rebound similar to the beta band, but with slower dynamics (Jurkiewicz et al. 2006; Pfurtscheller and Lopes Da Silva 1999; Donner et al. 2009; Engel and Fries 2010; Cheyne 2013; Barone and Rossiter 2021). It has been attributed a role in shaping the activation state of cortical sensory regions to affect the flow of information and optimize task performance (Jensen and Mazaheri 2010a; Haegens et al. 2011b; Spitzer and Blankenburg 2011; 2012). This could be mediated in part by regulating the level or direction of attention during decision-making (Vanni et al. 1997; Kelly et al. 2006; 2009; Foxe and Snyder 2011; Gould et al. 2011; Klimesch 2012; Van Diepen et al. 2019; Samaha et al. 2017; 2020).

Our key finding was a robust beta-band power increase shortly after auditory feedback that confirmed a correct decision, particularly in anterior cingulate, ventral and dorsal premotor and superior parietal regions, and some primary sensory areas. Critically, the feedback-evoked beta rebound magnitude was minimal in strong-evidence trials, when correct outcomes were expected. In contrast, it increased sharply when decisions were made under weak evidence, when experienced error rates were higher and correct outcomes were less likely. These evidence-dependent feedback-evoked response modulations are consistent with a positive RPE calculation. In contrast, the +RPE-like feedback-evoked response was weak in the primary motor (M1) and somatosensory (S1) cortex, indicating that those central sensorimotor regions do not contribute strongly to post-feedback aspects of performance monitoring and embodied decision-making in these tasks. We also observed RPE-like post-feedback responses in the alpha band, that were weaker and more spatially restricted to non-motor regions compared to the beta band.

In contrast, both alpha and beta power during the time between the perceptual-report hand movement and the reception of the outcome feedback signal scaled positively with sensory evidence strength or alternatively was suppressed while waiting for the outcome feedback in trials with Weak evidence. This anticipatory modulation may reflect internally computed evidence-dependent outcome expectations, another critical element in the RPE computation.

## Materials and Methods

### Participants

The study included 16 healthy participants (9 men and 7 women; mean age: 25.6 ± 3.6 years), all with no history of neurological or psychiatric disorders and no ferrous-metal implants. All participants were naïve to the objectives of the study. Two participants were ultimately excluded due to insufficient MEG signal quality, resulting in a final cohort of 14 participants. All 14 participants were right-handed, had normal or corrected-to-normal vision, no known color vision deficiencies, and were able to easily distinguish the colored cues used in the task. One author (J.K.) also participated. He was highly experienced with the tasks, but his behavioral performance and MEG data were within the group range. The study protocol was approved by the Institutional Review Board of the Montreal Neurological Institute, and all participants provided prior written informed consent.

All behavioral, MEG and anatomical MRI recordings were conducted in the McConnell Brain Imaging Centre, Montreal Neurological Institute, McGill University.

### General study design

Participants completed four variants of a color discrimination task (Supplemental Table 1). In each task, they judged the dominant color of a dynamic checkerboard-like stimulus (CKB) composed of different numbers of pale blue (B) and orange (O) squares against a neutral gray (G) background. Two monochromatic color-coded Target Cues—one B and one O—appeared on opposite sides of the CKB. Participants reported their perceptual decision by pressing either the left or right mouse button with the corresponding index finger, following a simple color-location conjunction rule: the spatial position of the selected button matched that of the Target Cue whose color corresponded to the perceived dominant color of the CKB. Participants either reported their decision as soon as they were ready (TFRT and CFRT tasks) or after an imposed delay (TFD and CFD tasks; Supplemental Table 1).

The strength of sensory evidence supporting the correct color choice was manipulated by varying the ratio of B and O squares across trials (see *Checkerboard Stimuli*). The left/right spatial positions of the B and O Target Cues were randomized on each trial, dissociating perceptual and motor components. The temporal order of CKB and Target Cue presentation differed across tasks to dissociate in time perceptual decision-making from motor preparation. These tasks were adapted from prior psychophysical studies in humans and neurophysiological recordings in NHPs (Coallier and Kalaska 2014; Coallier et al. 2015; Wang et al. 2019).

All participants completed three sessions on non-consecutive days within a two-week period (Figure 1A).

- **Sessions 1–2:** Reaction-time tasks—Target-First Reaction Time (TFRT; Figure 1B) and Checkerboard-First Reaction Time (CFRT; Figure 1C)—were performed in counterbalanced order across participants. Each task consisted of four consecutive trial blocks during which all experimental conditions were presented in pseudo-random order and served for training and baseline behavioral data collection.
- **Session 3:** Delayed-response versions of the same tasks—Target-First with Delay (TFD; Figure 1D) and Checkerboard-First with Delay (CFD; Figure 1E)—were performed during MEG recording. Each task likewise included four trial blocks. They were accompanied by separate resting-state and Random-Beep control blocks (Figure 1A).

**Figure 1.**
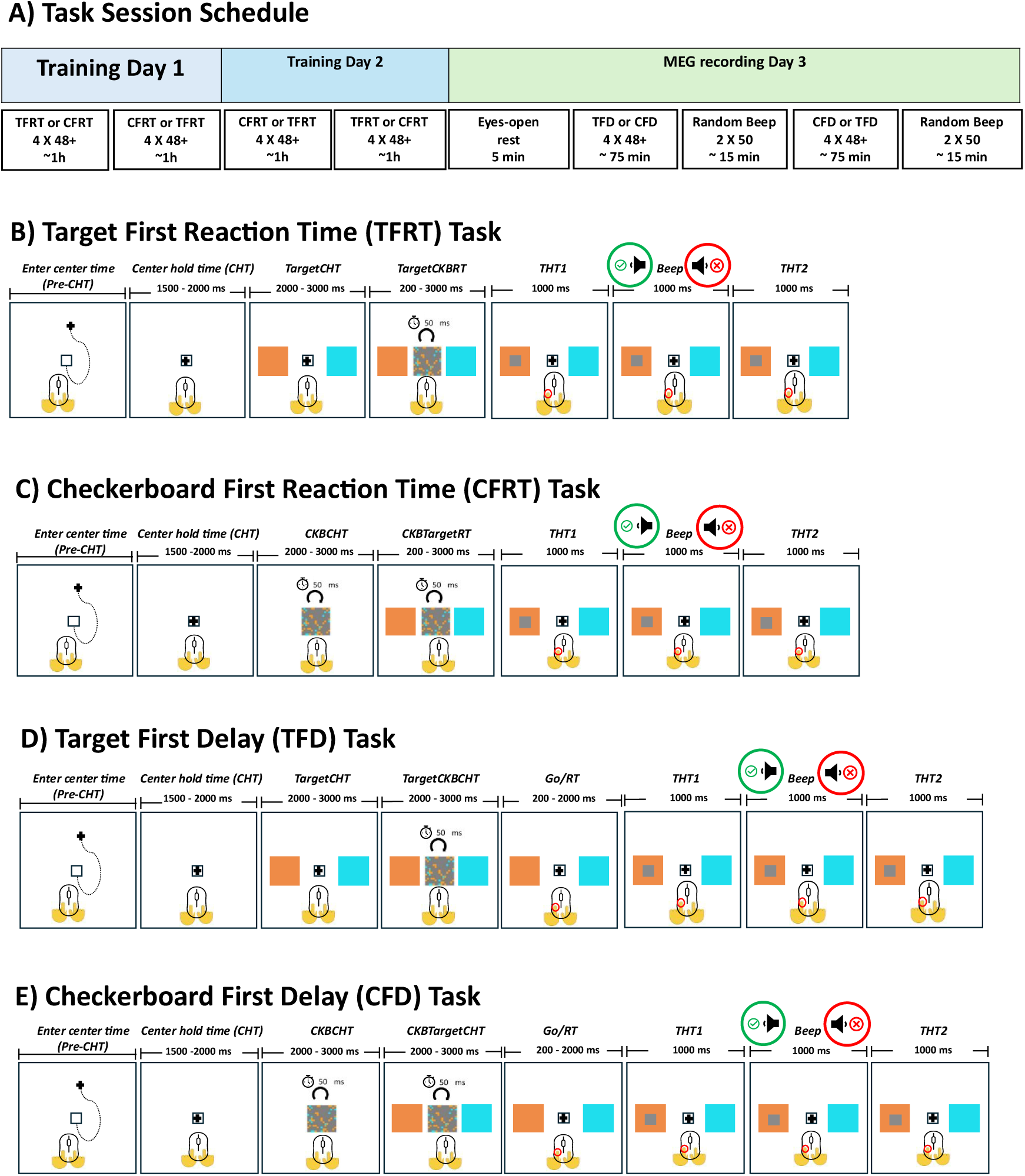
Overview of task structure and timeline across training and MEG sessions. **(A)** The study included three separate, non-consecutive daily sessions. On Training Day 1 and Training Day 2, participants performed reaction-time versions of two tasks (TFRT and CFRT) to become familiar with the procedures and to establish baseline behavioral performance. In these tasks, participants pressed a mouse button immediately upon making their perceptual and motor decisions—prompted by the appearance of the Checkerboard cue (CKB) in the TFRT task (TargetCKBRT epoch; panel **(B)**, or by the appearance of the Target Cues in the CFRT task (CKBTargetRT epoch; panel **(C)**. On Day 3, participants performed delayed-response versions of these tasks (TFD and CFD) while undergoing MEG recording (panels D and E). These tasks were nearly identical to the reaction-time versions, but with the addition of a second delay period after the CKB (TargetCKBCHT epoch; panel **(D)** or after the Target Cues (CKBTargetCHT epoch; panel **(E)**. In the delayed tasks, participants were required to withhold their report response until the disappearance of the CKB and the reappearance of the central fixation square, which signaled the start of the button-press response (Go/RT epoch; panels D and E).

Participants were allowed a self-timed rest period between each trial block in all tasks. They could also pause the task at any time during a trial block, but rarely did.

### Checkerboard stimuli

Each CKB consisted of a 5.7 cm square matrix (3.27° visual angle) composed of 15×15 smaller squares (0.38 cm) (Figure 2). Task-relevant squares were either isoluminant Blue (B) or Orange (O), and the remaining “background” squares were a neutral Gray (G). The exact CIELAB coordinates were O = [67.00, 42.63, 73.12], B = [67.00, –7.41, –50.92], and G = [53.39, 0, 0], selected after extensive psychophysical testing (Milosz 2021) to ensure that B and O were equidistant and oppositely located relative to G in CIELAB color space. They were converted to RGB for display: O = [254, 128, 6]; B = [3, 172, 254]; G = [128, 128, 128]. Because there were no visible borders between like-colored squares, adjacent squares of the same color appeared visually fused, forming irregularly-shaped clusters (Figure 2).

**Figure 2.**
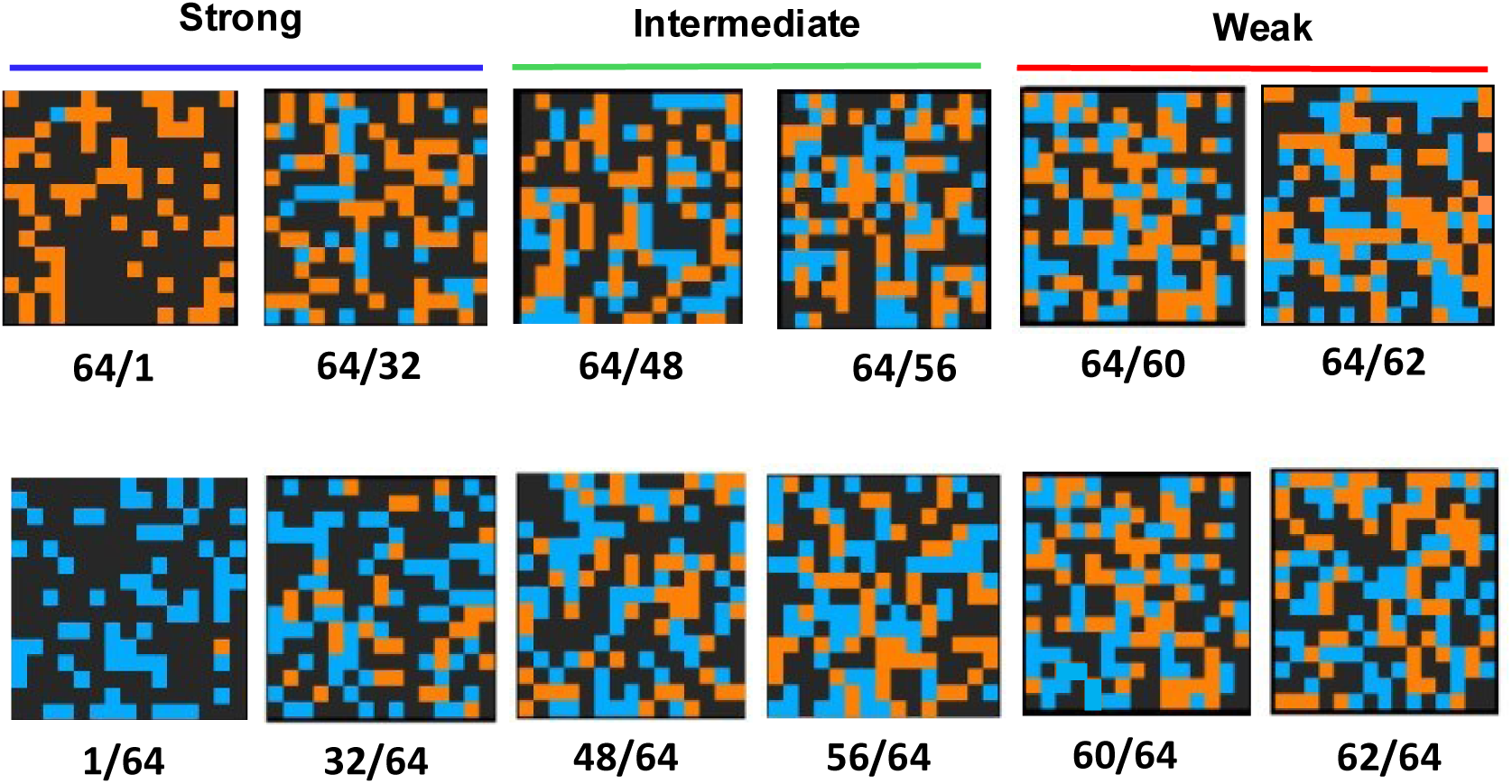
Examples of the range of individual checkerboard (CKB) matrices used in the task. The top row shows examples that were predominantly orange; the bottom row shows those that were predominantly blue. For each of the 12 combinations of conflicting color evidence strength, the task software included pre-generated libraries of 500 CKB matrices with randomized spatial arrangements of colored squares. In each trial, a random sequence of matrices from the appropriate library was presented at 50ms intervals. Note that the colors displayed here do not exactly match their true appearance in the original task displays.

Participants judged whether the CKB was predominantly blue or orange, that is, whether it contained more B or O squares. The number of squares of the dominant color (i.e., the correct color choice) was fixed at 64 in all CKBs. Relative sensory evidence strength was systematically manipulated by varying the number of squares of the incorrect color choice, ranging from 1, 32, 48, 56, 60, to 62 squares (Figure 2). This yielded twelve unique CKB color combinations (six B-dominant and six O-dominant). As the number of incorrect-color squares approached that of the correct color, sensory evidence for the two opposite choices became more balanced, increasing decision difficulty.

For analytical purposes, the 6 levels of evidence strength were grouped into three categories of net evidence strength supporting the correct B or O choice, **Strong:** (64/1, 64/32); **Intermediate:** (64/48, 64/56); and **Weak:** (64/60, 64/62). Quantitatively, the mean net evidence supporting the correct color choice was 47.5 squares for Strong CKBs (64−[(1+32)/2]), 12 squares for Intermediate CKBs, and 3 squares for Weak CKBs.

To prevent counting strategies, each color ratio condition was presented as a dynamic sequence of CKBs with randomized B and O square locations in each trial (Coallier & Kalaska, 2014; Coallier et al., 2015). For every ratio, a library of 500 unique CKB matrices was created, with randomly distributed B and O squares. Each matrix had a complementary version with reversed colors (e.g., 64B/32O ↔ 64O/32B), ensuring spatial counterbalancing between color-dominant conditions. During each trial, a random sequence of matrices was drawn from the desired color ratio library and presented in rapid succession every 50 ms. This maintained a constant color ratio in a trial while continuously changing the spatial distribution of B and O squares within the CKB. This minimized local luminance or positional biases and emphasized global perceptual evaluation of color dominance.

### Target-First Reaction Time (TFRT) task

Each TFRT trial began with the appearance of an open white square (1.9 cm) at the center of the screen (Figure 1B; Supplemental Table 1). Participants placed the index fingers of their left and right hands on the respective buttons of a computer mouse positioned on a flat surface at waist level. They moved the mouse to align the cursor within the central square, marking the end of the enter-center epoch (Pre-CHT) and the beginning of the center hold time (CHT). Participants typically maintained the cursor in the central square throughout the trial block. The CHT lasted 1500– 2000 ms, randomized across trials. Two peripheral 5.7 cm square Target Cues appeared next, positioned 13.5 cm apart on either side of the central square, initiating the TargetCHT epoch (2000–3000 ms, randomized). The Target Cues were color-coded (one Blue, one Orange), with their left-right positions randomized across trials.

Subsequently, the central square was replaced by a 5.7 cm dynamic CKB containing randomly arranged B, O, and G squares. The CKB onset initiated the TargetCKBRT epoch. Participants evaluated the dominant color in the CKB and selected the matching Target Cue by pressing the corresponding left or right mouse button, according to the predefined color-location conjunction rule.

Participants were free to make a report mouse-button press as soon as they were ready after CKB onset at the start the TargetCKBRT epoch (valid RTs 200–3000 ms). Once a button press was detected, the CKB disappeared to prevent post-decision viewing and to minimize potential “Change of Mind” responses (Burk et al. 2014; Coallier and Kalaska 2014; Dotan et al. 2019; Resulaj et al. 2009). Simultaneously, a darker 1 cm square (B: [0,130,220]; O: [220,130,0]) appeared at the center of the chosen Target Cue to mark the response (shown in gray in Figure 1B–E). Target Cues remained visible until trial end. Participants had to continue pressing the selected mouse button during the Target Hold 1 epoch (THT1; 1000ms, fixed duration). At the end of THT1, a 1000ms auditory tone (BEEP epoch) was delivered via earphones. The Feedback-Beep cues provided feedback on decision accuracy: a 1500 Hz tone indicated a correct decision, and a 250 Hz tone an incorrect one. After Feedback-Beep offset, participants held the mouse button for an additional 1000ms during the Target Hold 2 epoch (THT2). After THT2, all visual stimuli disappeared, and an identical green message appeared near the top of the screen for 1 second: “Trial Over – Release the Button!”, regardless of trial outcome. The auditory Feedback-Beep tones were the sole indicator of decision outcome (correct or incorrect). Each trial was followed by a 2500ms inter-trial interval.

Report mouse-button presses during the TargetCKBRT epoch were self-paced, with no explicit emphasis on speed or accuracy. Participants were instructed to fixe their gaze centrally and avoid blinks during trials. Eye movements and blinks were allowed during inter-trial intervals. However, no gaze monitoring or gaze enforcement was applied.

### Checkerboard-First Reaction Time (CFRT) task

The CFRT task reversed the cue order used in TFRT to separate perceptual evaluation from motor preparation (Figure 1C; Supplemental Table 1; Coallier and Kalaska 2014; Coallier et al. 2015; Wang et al. 2019). Following the initial CHT period, the CKB appeared first and remained visible for 2000–3000 ms (CKBCHT epoch). Participants could estimate the dominant color without yet knowing the color-location mapping, as the Target Cues had not been presented. Subsequently, the color-coded Target Cues appeared to initiate the CKBTargetRT epoch. This served as the instructional cue enabling participants to apply the conjunction rule: select the Target Cue whose color matched the estimated dominant color of the CKB and report the decision by pressing the corresponding left or right mouse button, based on the spatial position of the selected Target Cue (valid RTs 200–3000 ms). All subsequent epochs (THT1, BEEP, THT2) and timing parameters were identical to the TFRT task. Apart from the reversed cue sequence, all visual stimuli, timing, and counterbalancing procedures were identical across the two reaction-time tasks.

### MEG recording session

In the third, non-consecutive daily session, participants completed four trial blocks of instructed-delay versions of the two tasks: Checkerboard-First with Delay (CFD) and Target-First with Delay (TFD). Each participant performed both tasks during the MEG recording session, with task order counterbalanced across participants. Additional short MEG acquisition blocks were also included (Figure 1A). Visual stimuli were presented via back-projection onto a translucent screen positioned at eye level, approximately 1 meter in front of the participant seated in the MEG chair. Stimuli were displayed using a PROPixx projector (VPixx Technologies) with a 120-Hz frame refresh rate (https://vpixx.com/).

### Target-First with Delay (TFD) task

The TFD task was a modified version of the TFRT task adapted for MEG recording (Figure 1D; Supplemental Table 1). Each trial in the TFD task followed the same sequence of epochs as in TFRT up to the presentation of the CKB. However, unlike TFRT, participants were not permitted to press a mouse button as soon as they had decided on the dominant color of the CKB after it appeared.

Instead, an additional delay period was imposed after CKB onset (TargetCKBCHT epoch; 2000–3000ms, randomized), during which participants had to withhold their button-press response. The end of this delay was marked by the disappearance of the CKB and reappearance of the central white square, serving as the “Go” cue. Participants then reported their decision with a button press as quickly as possible during the Go/RT epoch (valid RTs 200–2000ms). The remainder of the trial (THT1, BEEP, and THT2 epochs) was identical to the TFRT task.

### Checkerboard-First with Delay (CFD) task

The CFD task was the delayed-response version of CFRT (Figure 1E; Supplemental Table 1). Its structure was identical to CFRT except for an added delay period following Target Cue presentation. After the Target Cues appeared to start the CKBTargetCHT epoch (2000–3000 ms, randomized), participants withheld their mouse button press. The Go/RT epoch began when the CKB disappeared and the central white square reappeared, signaling participants to make their button-press response (valid RTs 200–2000ms). The remainder of the trial, comprising the THT1, BEEP, and THT2 epochs, proceeded exactly as in the CFRT task.

The purpose of the additional delay in the TFD and CFD tasks was to dissociate in time the neural processes underlying perceptual and motor decision-making from those associated with the physical execution of the motor response. The visual stimuli used in the TargetCKBRT, TargetCKBCHT, CKBTargetRT, and CKBTargetCHT epochs were identical across all four task variants. However, differences in cue order and response constraints were expected to yield distinct neural dynamics. For clarity, distinct epoch labels were maintained for each task.

### Random-Beep trials

Following the completion of each TFD and CFD task on Day 3, participants performed a passive auditory listening task referred to as the Random-Beep condition. During this task, participants sat quietly while maintaining their gaze on a central white square displayed on the screen. They rested their index fingers on the left and right mouse buttons but made no responses. The Random-Beep condition measured neural responses to the auditory feedback tones in a task-free context that did not require decision outcome-feedback processing.

Each Random-Beep block consisted of a randomized sequence of 50 “correct” tones (1500 Hz) and 50 “incorrect” tones (250 Hz), each lasting 1000ms, that were physically identical to the Feedback-Beep signals delivered during the TFD and CFD tasks. Tones were presented at an average interval of 5000ms. One block was completed at the end of each delayed-response task (TFD and CFD), yielding two Random-Beep blocks and a total of 100 “correct” trials and 100 “incorrect” trials per participant.

### Trial block structure

Each task comprised six levels of sensory evidence, two possible dominant colors (B or O), and two correct target spatial locations, yielding 24 unique trial conditions. Within each trial block, participants had to perform all 24 trial conditions correctly two times, for a total of 48 correct trials per block. Trial order was pseudo-randomized to prevent sequence predictability. A trial was considered **correct** if the participant pressed the mouse button corresponding to the Target Cue whose color matched the dominant color of the CKB and was on the appropriate side according to the color–location conjunction rule, and completed the rest of the trial successfully until the end of the THT2 epoch. A trial was considered **incorrect** if the participant selected the Target Cue whose color was inconsistent with the CKB’s dominant color, but completed the rest of the trial. If a participant selected the incorrect target, the data for that trial were stored with a unique error identifier and the trial condition was reinserted into the trial sequence to be repeated later until performed correctly.

Task performance was also monitored for other error conditions, including premature mouse button presses during the delay periods before the TargetCKBRT (TFRT), CKBTargetRT (CKBRT) and Go/RT (TFD, CFD) epochs, RTs that were too short or too long, and premature mouse button release during the THT1, Beep or THT2 epochs. If any of these performance errors were detected, the trial was terminated immediately, and the data were stored with unique error type identifiers. A red text message also appeared at the top of the display to inform the participant of the nature of the error. The trial condition for those performance errors was also reinserted into the trial sequence to be repeated later. These performance errors were rare and will not be described in any further detail in this report.

These trial management procedures ensured that each block included exactly 48 completed **correct** trials, and variable numbers of completed **incorrect** trials, and other performance error trials. Each participant completed 4 blocks per task, totaling 192 correct trials per task, fully counterbalanced across all trial conditions.

### MEG and structural MRI data collection

The MEG recording session was conducted using a 275-channel VSM/CTF system (https://www.ctf.com/) located in a magnetically shielded room with three layers of passive shielding to minimize environmental interference. Data were sampled at 2,400 Hz, with a hardware anti-aliasing low-pass filter applied at 600 Hz to reduce high-frequency noise and ensure data fidelity.

To monitor eye movements and minimize ocular artifacts, electro-oculogram (EOG) electrodes were placed at the outer canthi and above and below the left eye, capturing both horizontal (HEOG) and vertical (VEOG) movements, as well as blinks and large saccades. An electrocardiogram (ECG) was also recorded to identify cardiac-related artifacts, using electrodes placed across the chest below the clavicles. All auxiliary signals were recorded at the same sampling frequency as the MEG data.

Auditory feedback tones were delivered using E-A-RTone insert earphones fitted with disposable foam tips. The default sampling rate of the audio system was 44,100 Hz. This comprehensive acquisition setup ensured optimal conditions for the collection of high-quality MEG data suitable for sensor- and source-level analyses.

For MEG source mapping, participants also underwent a T1-weighted structural MRI scan (3T Siemens Prisma Trio), either on the same day as the MEG session or on a separate day. Co-registration of MRI and MEG data was achieved using three fiducial points (nasion, left and right pre-auricular) digitized with a Polhemus 3-D digitizer (Isotrak, Polhemus Inc., VT, USA) at the start of the MEG session. This alignment was refined using approximately 100 additional digitized scalp points processed through Brainstorm. Cortical and scalp surfaces were generated from individual MRIs using FreeSurfer (default parameters) and down-sampled to 15,002 cortical vertices for MEG source modeling.

During MEG recordings, participants sat upright in a comfortable chair and were instructed to fixate on a centrally presented task display at eye level. The MEG recording session began with a minimum of five minutes of resting-state recording with eyes open (Figure 1A). During this period, participants maintained steady gaze on a fixation cross. These resting-state data are part of The Open MEG Archive (OMEGA), a publicly accessible repository supporting future research into brain function (Niso et al. 2019).

Following the resting-state data acquisition, participants performed both the TFD and CFD tasks, with task order counterbalanced across participants to mitigate potential order effects. Each decision-making task was followed by a passive auditory control block of 100 Random-Beep trials (Figure 1A).

### MEG data preprocessing

All MEG preprocessing and source imaging were performed using Brainstorm (Tadel et al. 2011), supplemented with custom MATLAB scripts (R2023b, The MathWorks Inc., Natick, MA, USA). Full implementation details are publicly documented and can be verified in the Brainstorm code repository. MEG recordings incorporated the built-in third-order gradient compensation provided by the CTF system.

To correct for variable delays between stimulus presentation commands and actual visual display onset due to projector latency, a photodiode was used in the MEG room to register the timing of each visual stimulus in every trial. Power line noise (60 Hz and its harmonics) was attenuated using 3-dB bandwidth notch filters.

Bad sensor channels were identified via visual inspection of their power spectra and flagged for exclusion. Artifacts caused by cardiac activity, eye blinks, saccades, and muscle contractions were addressed using signal-space projection (SSP) techniques, following recommended best practices (Gross et al. 2013). Contaminated epochs were automatically detected based on signal variance within specific frequency bands—for example, 1.5–15 Hz for eye blinks and 10–40 Hz for cardiac artifacts. SSPs were computed from time windows centered on the artifact events (typically 400ms around eye blinks and 160ms around heartbeats). Remaining artifacts were removed by manual review, ensuring high-quality data for sensor- and source-level analyses.

Time stamps stored at the start of each trial epoch during MEG data collection were used to segment the MEG data into defined time windows that typically spanned −2000 ms to +3000 ms relative to each trial epoch onset. Time windows shorter than 5 s were excluded. Artifact rejection used a ±3 MAD threshold on both amplitude and gradient at the single-trial level, following Wiesman et al. (2022). Only artifact-free epochs were retained for analysis.

### Behavioral analysis -***reaction times and error rates

Behavioral data from both the training and MEG recording sessions were analyzed using custom MATLAB scripts. Only trials in which participants completed all trial epochs without committing a performance error and selected either the correct or incorrect target were analyzed. Trials involving all other types of performance errors (e.g., anticipations, guesses, premature releases) were excluded.

**Error rate** was defined as the percentage of incorrect target selections relative to the total number of correct and incorrect trials at each level of sensory evidence strength. This metric was used to assess decision accuracy as a function of the net color evidence strength of the CKB cues, reflecting how well participants could identify the dominant color under varying levels of evidence ambiguity.

**Reaction time (RT)** was measured as the interval between the onset of the decision-report epoch—TargetCKBRT (TFRT), CKBTargetRT (CFRT), or Go/RT (TFD and CFD)—and the detected onset of the participant’s mouse button press. RT served as an indirect measure of the time required for perceptual evaluation, decision-making, and motor response preparation and execution in different task conditions.

### MEG Source Imaging

Source estimates for each participant were re-projected onto a common template brain to facilitate inter-subject comparisons. Specifically, we used the ICBM152 cortical anatomy with 15,002 vertices as the standard source space. Each participant’s source reconstruction was aligned to this common space via registration to the FSaverage spherical template, allowing vertex-wise correspondence across individuals. This enabled group-level averaging and statistical analyses of source-level MEG data across the cohort.

### Time-Frequency Analysis of MEG Oscillatory Activity

We performed frequency-resolved analyses of the MEG source signals at each cortical vertex location. For each trial, source signals were segmented into temporally overlapping sequential time windows spanning −2000ms to +3000ms relative to the onset of each trial epoch.

Each single-trial time window was band-pass filtered into canonical frequency bands using an even-order linear phase finite impulse response (FIR) filter with 60 dB stopband attenuation. The frequency bands included: delta (2–4 Hz), theta (5–7 Hz), alpha (8–12 Hz), beta (15–29 Hz), and low gamma (30–59 Hz). The Hilbert transform was then applied to the filtered signals to derive the analytic signal, from which the instantaneous amplitude (envelope) and phase were computed for each time point and vertex.

To reduce boundary artifacts from the filtering and Hilbert transform operations, all subsequent analyses were confined to the central portion of each time window, from −1000ms to +2000ms relative to trial epoch onset. A common baseline was defined as the −1000ms to 0ms interval preceding the start of the Center Hold Time (CHT) epoch at the beginning of each trial.

Single-trial data were first categorized into Strong, Intermediate, and Weak evidence conditions. Within each category, the Hilbert transform was applied to the single-trial source-level brain maps, which were then averaged across all trials. The regional mean responses were subsequently computed by averaging across all vertices within each region of interest (ROI) separately for the left and right hemispheres, followed by averaging across the bilaterally matched ROIs. Finally, the resulting mean temporal profiles in each ROI were averaged across participants.

### Regions of interest (ROIs)

We analyzed MEG signals in a targeted set of a priori defined anatomical/functional regions of interest (ROIs; Figure 3), selected based on extensive literature describing the organization of the human cortical sensorimotor system (e.g., Badre 2008; Binkofski and Buccino 2006; Chouinard and Paus 2006; 2010; Rizzolatti et al. 2002). These included bilateral hand/arm representations in the primary motor cortex (M1) and primary somatosensory cortex (S1), the dorsal (PMd) and ventral (PMv) premotor cortex, and the supplementary and pre-supplementary motor areas (preSMA/SMA). The PMv ROI encompassed the lateral portion of Brodmann area 6 and the caudal portion of neighboring area 44 (a part of Broca’s area).

**Figure 3.**
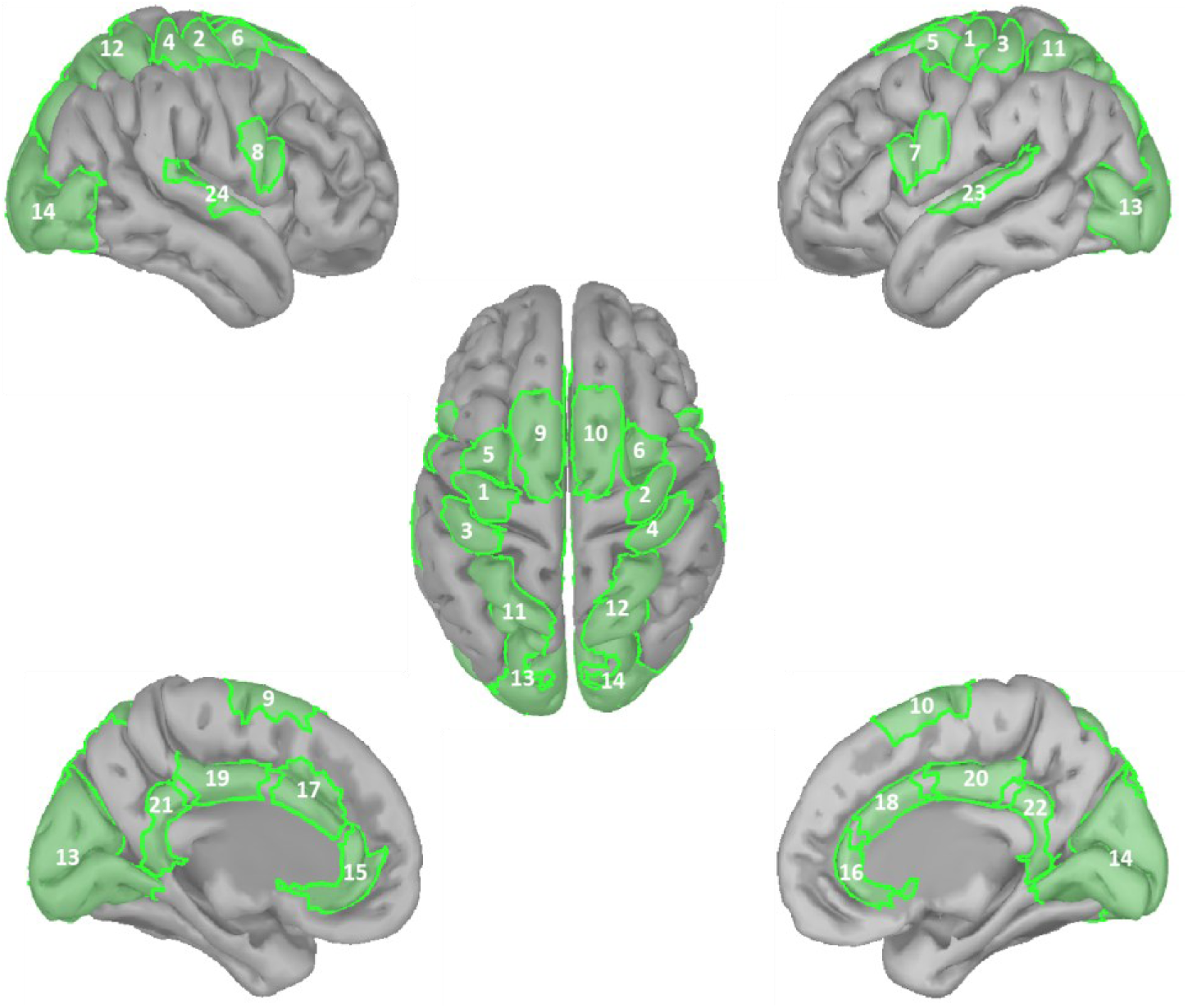
Bilaterally symmetric regions of interest (ROIs) used in analyses. The defined ROIs include: 1–2) Primary motor cortex (M1, hand/arm region); 3–4) Primary somatosensory cortex (S1, hand/arm region); 5–6) Dorsal premotor cortex (PMd); 7–8) Ventral premotor cortex (PMv; ventral BA 6 and 44); 9–10) Supplementary motor area and pre-supplementary motor area (SMA/preSMA); 11–12) Superior parietal lobule (SPL; BA 5 and 7); 13–14) Occipital pole (BA 17, 18, 19); 15–16) Rostral anterior cingulate cortex; 17–18) Caudal anterior cingulate cortex; 19–20) Posterior cingulate cortex; 21–22) Isthmus cingulate cortex; 23–24) Primary auditory cortex.

We also included the rostral anterior cingulate, caudal anterior cingulate, posterior cingulate and isthmus cingulate cortex and the superior parietal cortex, as defined by the Desikan-Killiany-Tourville (DKT) atlas (Desikan et al. 2006). To capture MEG activity in early visual and auditory sensory cortices, we defined the occipital pole (Brodmann areas 17, 18, and 19) as a visual ROI, and we used the Schaefer-100 parcels SomMotB Aud 1L and SomMotB Aud 1R as an auditory ROI (Schaefer et al. 2018).

### Quantification of MEG signals evoked by auditory feedback signals about trial outcome

In this study, we focused on the MEG activity from the detected onset of the motor-report button press to the end of the trial, encompassing the THT1, Beep, and THT2 epochs (Figure 1), with emphasis on the signals evoked by the auditory Feedback-Beep signals. We also focussed here on trials with correct outcomes. MEG signals asociated with incorrect decisions, and activity generated during trial epochs preceding the the mouse-button motor response will be reported separately.

Analyses focused on how sensory color evidence strength modulated feedback-evoked MEG responses in correct trials only, pooling across correct color (B or O) and target laterality (hemisphere contralateral or ipsilateral to the chosen mouse button). Correct trials were grouped by evidence strength: **Strong** (64/1, 64/32), **Intermediate** (64/48, 64/52), and **Weak** (64/60, 64/62). To assess the statistical significance of the effect of color evidence strength on the magnitude of the measured MEG responses, we used permutation tests for dependent (i.e., within-subject) samples (*permutest*, MATLAB R2023b) to compare distributions across task conditions within ROIs, and one-sample t-tests (*ttest*, MATLAB R2023b) on the pairwise differences in activity between trial groups pooled across ROIs.

Post-decision MEG data were analyzed separately from both TFD and CFD tasks. However, no task-specific effects were expected a priori on post-decision activity. Indeed, permutation tests revealed no significant post-decision differences between TFD and CFD data across ROIs at any evidence level (p > 0.05, FDR-corrected). Consequently, post-decision MEG responses from both tasks were pooled for all subsequent analyses.

### Beta band activity

Figure 4A illustrates the typical temporal dynamics of beta-band power changes (mean ± s.e.m.) observed across most ROIs during the sequential THT1, Beep, and THT2 epochs in correct trials. Consistent with many previous reports of movement-related beta band responses (Jurkiewicz et al. 2006; Pfurtscheller and Lopes da Silva 1999; Engel and Fries 2010; Cheyne 2013; Barone and Rossiter 2021; Rogge et al. 2022; An et al. 2019; Chouinard and Paus 2006; Kilavik et al. 2013; Tan et al. 2016), beta power exhibited progressive suppression below baseline in the trial epochs prior to the decision-report mouse button press (not shown), reaching a minimum at the time of the motor response that initiated the THT1 epoch (Figure 4A, time -1sec). This suppression was followed by a rapid post-report-movement beta rebound that peaked during the THT1 epoch and then declined sharply before the onset of the Beep epoch (Figure 4A, time 0sec). Importantly, however, shortly after the presentation of the auditory Feedback-Beep signal, a second beta rebound emerged, that peaked during the Beep epoch and then declined gradually throughout the remainder of the Beep and THT2 epochs. The feedback-evoked beta rebound component was the main focus of this study.

**Figure 4.**
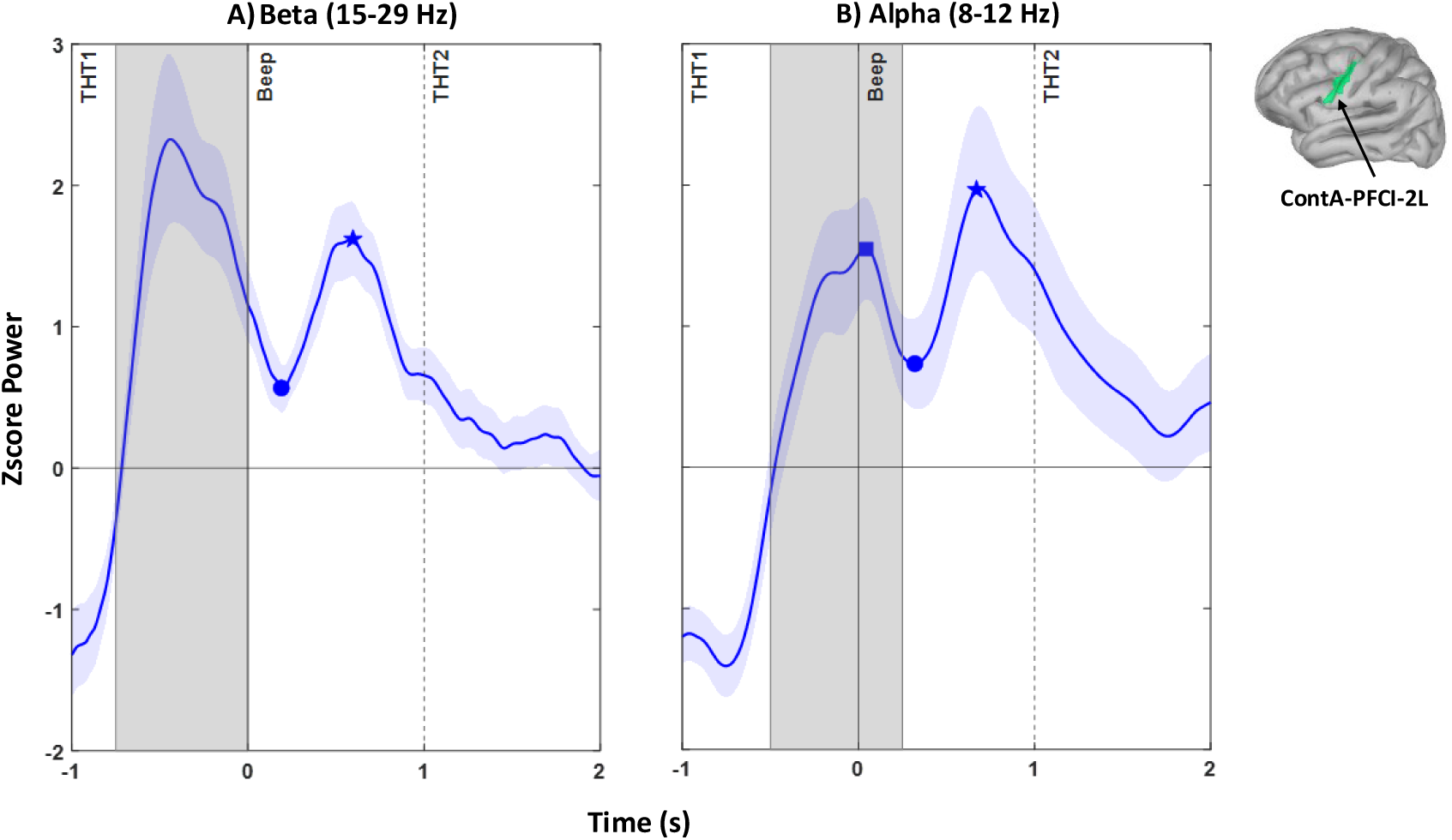
Typical temporal dynamics of beta-band and alpha-band power observed across many cortical regions during the post-decision time window of interest in correct trials. (A) Grand mean beta-band power sourced to Schaefer region ContA PCFl 2L, which includes parts of lateral Brodmann area 6 in the precentral gyrus and adjacent area 44 in the pars opercularis of the inferior frontal gyrus in the left hemisphere (see inset). Data are from the CFD task in trials with Strong color evidence (64/1 and 64/32 CKBs), pooled across B-dominant and O-dominant CKBs and across contralateral and ipsilateral target choices. The button press that initiated the THT1 epoch occurred at time –1 s; the onset of the Beep epoch is indicated by the solid vertical line at 0 s, and the onset of the THT2 epoch by the dashed vertical line at 1 s. (B) Grand mean alpha-band power from the same region (ContA PCFl 2L), task (CFD), and trial subset (strong color evidence), pooled using the same conventions as in (A). Same display format as in (A).

Three beta-band measures were quantified in each bilateral ROI and evidence condition. (1) The initial post-press rebound magnitude was computed as the mean z-scored beta power from −750 to 0 ms relative to Beep onset (Figure 4A, gray patch). (2) The subsequent local minimum was identified between −100 and +1000 ms (Figure 4A, circle; *findpeaks*, MATLAB R2023b). (3) The Feedback-Beep–evoked rebound peak was located between +500 and +1500 ms (Figure 4A, star). The Feedback-Beep evoked beta rebound magnitude was defined as the z-scored power difference between this peak (star) and the preceding local minimum (circle).

### Alpha band activity

Figure 4B depicts typical alpha-band power dynamics during the post-decision period in correct trials that resembled many previous descriptions of alpha-band (8-12Hz) and mu-band (8-14Hz) responses in motor and sensorimotor decision-making tasks (Jurkiewicz et al. 2006; Pfurtscheller and Lopes da Silva 1999; Haegens et al. 2011a,b; Cheyne 2013; Barone and Rossiter 2021; Rogge et al. 2022). Alpha power decreased in the trial epochs preceding the decision-report button press (not shown), but unlike the beta band, alpha band activity continued to decrease after the motor-report button press, reaching a minimum 200-300ms after the start of THT1 (Figure 4B, time −1 s) This was followed by a rebound that was generally slower and less pronounced than in the beta band, and that peaked shortly after the onset of the auditory Feedback-Beep (time 0 s). The onset of the Feedback-Beep cue evoked a brief suppression and a second rebound that extended into the post-Beep THT2 epoch.

The initial post-button-press rebound was quantified by averaging z-scored alpha power from −500 ms to +250 ms relative to Beep onset (Figure 4B, gray patch). We identified the initial rebound peak (−100 to +1000 ms; square), subsequent minimum (+100 to +1000 ms; circle), and later Feedback-Beep-evoked rebound peak (+100 to +1500 ms; star). We calculated two response magnitude measures: (1) the depth of the transient suppression as the difference in z-scored power between the initial rebound peak (square) and the following local minimum (circle), and (2) the magnitude of the Feedback-Beep-evoked alpha rebound as the difference between the local minimum (circle) and the subsequent peak (star).

All time windows for the beta-band and alpha-band temporal profiles were empirically optimized to identify the desired oscillatory components across ROIs and participants as reliably as possible. However, visual examination of each mean response profile for each trial subset in each task and individual indicated that the alpha-band response profiles were more complex and variable than the beta-band profiles, which introduced a higher degree of measurement variability for the local minima and maxima in the alpha band data.

Finally, the means of the beta and alpha band response component magnitudes measured from the response profiles of individual trial subsets in individual participants as described above will always be larger than the apparent magnitude of the corresponding response components in the grand mean response profiles averaged across all participants, because of the idiosynchratic variabilty of the timing of local maxima and mimina in the averaged response profiles of individual participants.

## Results

### Behavioral results: Error rates

Participants decided whether the Checkerboard cue (CKB) was predominantly blue (B) or orange (O), and reported their perceptual decision by pressing the left or right mouse button with the corresponding index finger to indicate the location of the Target Cue that matched the chosen dominant color. The accuracy of their sensorimotor decisions was strongly modulated by the net color evidence in each CKB.

Error rate was defined as the number of incorrect trials divided by the total number of correct and incorrect trials for each CKB composition, collapsed across L and R target choices (Figure 5, left). For MEG analyses, trials were grouped into three evidence strength levels based on net color composition: Strong (64/1, 64/32), Intermediate (64/48, 64/56), and Weak (64/60, 64/62), collapsed across L/R targets and predominantly B and O CKBs (Figure 5, right).

**Figure 5.**
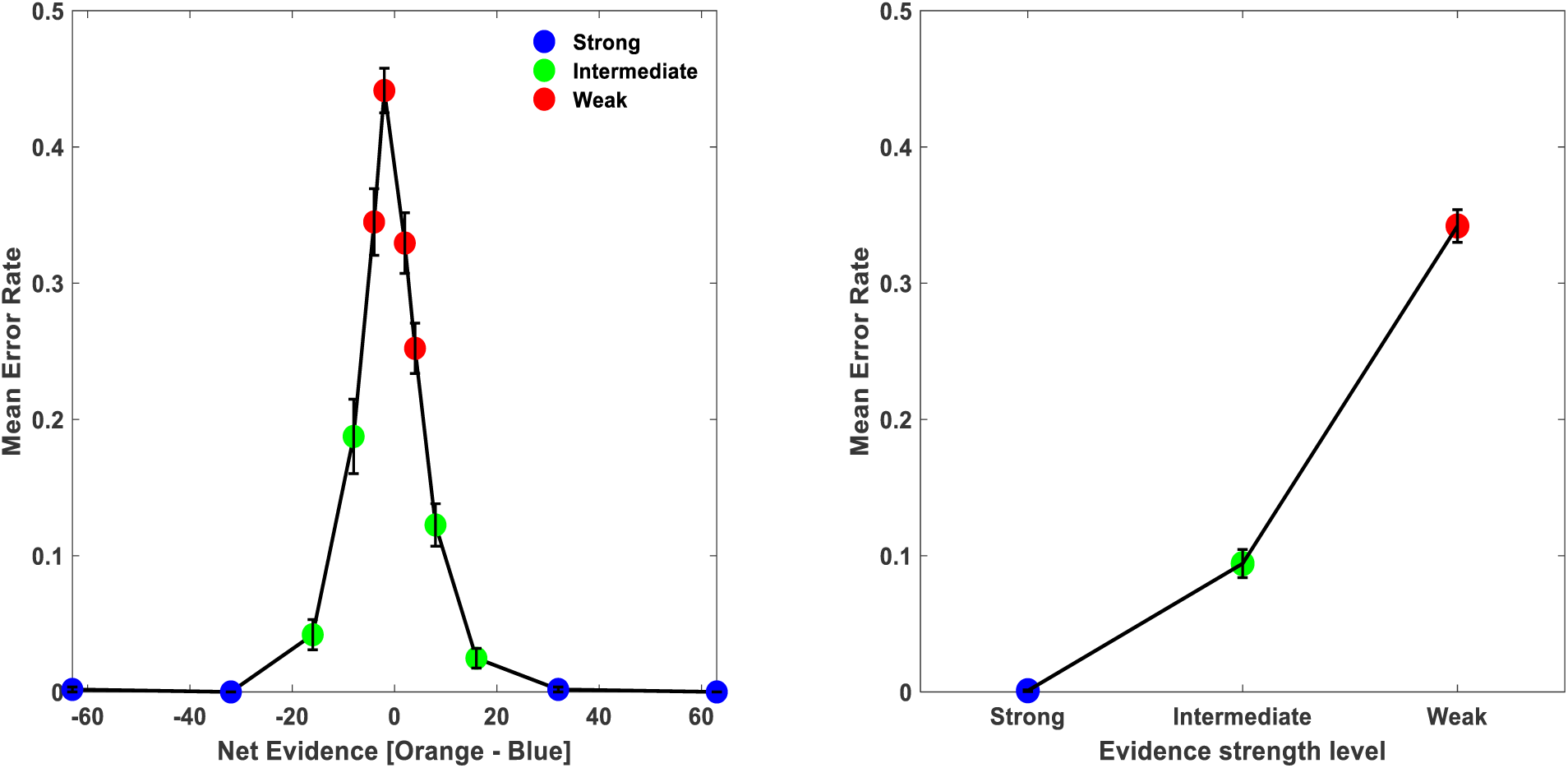
Grand mean error rates (± s.e.m.), first pooled across the CFD and TFD tasks within each participant and then averaged across all participants. Left panel: Error rates for all combinations of predominantly blue (B) and predominantly orange (O) checkerboard cues, pooled across L and R Target choices. Right panel: Error rates for trials grouped by the level of net color evidence for the correct decision, regardless of dominant color. Blue dots: Strong evidence trials (64/1, 64/32); Green dots: Intermediate evidence trials (64/48, 64/56); Red dots: Weak evidence trials (64/60, 64/62).

Error rates increased systematically as net color evidence decreased (Figure 5), consistent with previous studies using similar CKB stimuli (Coallier and Kalaska 2014; Coallier et al. 2015; Wang et al. 2019). Participants almost never made an incorrect decision in trials with Strong color evidence (mean error rate: 0.09% ± 0.06% s.e.m.; Figure 5, right, blue dot). Error rates increased progressively in trials with Intermediate (9.42% ± 1.03% s.e.m.; green dot) and Weak evidence (34.20% ± 1.19% s.e.m.; red dot). Nevertheless, even with the weakest CKBs (62/64 and 64/62), participants performed significantly better than chance (mean error rates: 44.14% ± 1.64% s.e.m. and 34.92% ± 2.45% s.e.m., respectively), indicating that perceptual decisions were informed by the available visual evidence in those trials.

Post-experiment debriefing revealed that participants were consciously aware of this performance gradient. All reported recognizing that they were highly unlikely to make an error in trials with Strong evidence but often made errors in trials with Weak evidence. Indeed, some described feeling they were merely guessing in trials with the weakest CKBs, despite mean success rates of ∼60.5%. This indicated that the color evidence strength not only had a systematic effect on decision accuracy but also had a systematic effect on the participants’ introspective trial-to-trial metacognitive awareness of their level of certainty about the accuracy of their decisions while waiting to receive auditory feedback about their decision outcomes after they pressed their chosen mouse button. However, because we did not ask participants to report their level of decision confidence or of their expectation of correct versus incorrect outcomes on a trial-by-trial basis, we cannot quantify their subjective outcome expectations relative to their objective observed outcome performance (Hajcak et al 2005, 2007; Ichikawa et al 2010; Hoy et al 2021).

The level of net color evidence in the CKBs and the order of presentation of the CKBs and Target Cues both also had a strong effect on report-movement RTs in the CFRT and TFRT tasks during training that indicated that processing of the color information in the CKBs accounted for most of the decision time in the tasks (Supplemental Figure 1A). This effect largely disappeared in the CFD and TFD tasks during MEG recordings (Supplemental Figure 1B), indicating that the participants had processed the color information provided by the CKBs and colored Target Cues and were committed to a button-press decision before the Go cue appeared at the end of the second delay period in both the TFD and CFD tasks. These evidence- and task-dependent RT modulations are consistent with previous reports (Coallier and Kalaska 2014; Coallier et al. 2015; Wang et al. 2019).

### “Correct” Random-Beep tone responses

After completing each of the TFD and CFD tasks during the MEG recording session, participants passively listened to a block of Random-Beep trials comprised of a randomized sequence of 50 “correct” (1500 Hz) and 50 “incorrect” (250 Hz) auditory tones (1000ms duration each) while fixating their gaze on a central white square on the screen. They placed their left and right index fingers on the left and right mouse buttons, but did not press a button. These “Random-Beep” trials were designed to assess brain responses to the tones used as the tasks’ outcome feedback signals when presented in a neutral context outside the task. Importantly, however, the Random-Beep condition was not intended to serve as a neutral “passive sensory” baseline that isolated pure sensory components of the feedback responses. Participants had already learned the association between tone frequency and correct versus incorrect outcomes during the prior training sessions, potentially imbuing the tones with an acquired implicit valence meaning even when presented outside of the decision-making tasks. As such, responses to Random-Beep tones likely reflect a mixture of sensory processing and an automatic recall of the learned valence meaning of the two auditory tones, but lack the evidence-specific and decision-specific contextual information present during task-embedded outcome feedback processing. Consequently, comparisons between Random-Beep and Feedback-Beep responses are informative about task specificity and contextual modulation, but not about purely sensory versus purely task-outcome evaluation processes.

Figures 6 and 7 illustrate the changes in grand mean z-normalized power in the beta band and alpha band, respectively, evoked by “correct” 1500Hz Random-Beep tones in the 12 a priori defined bilateral ROIs (c.f., Figure 3).

**Figure 6.**
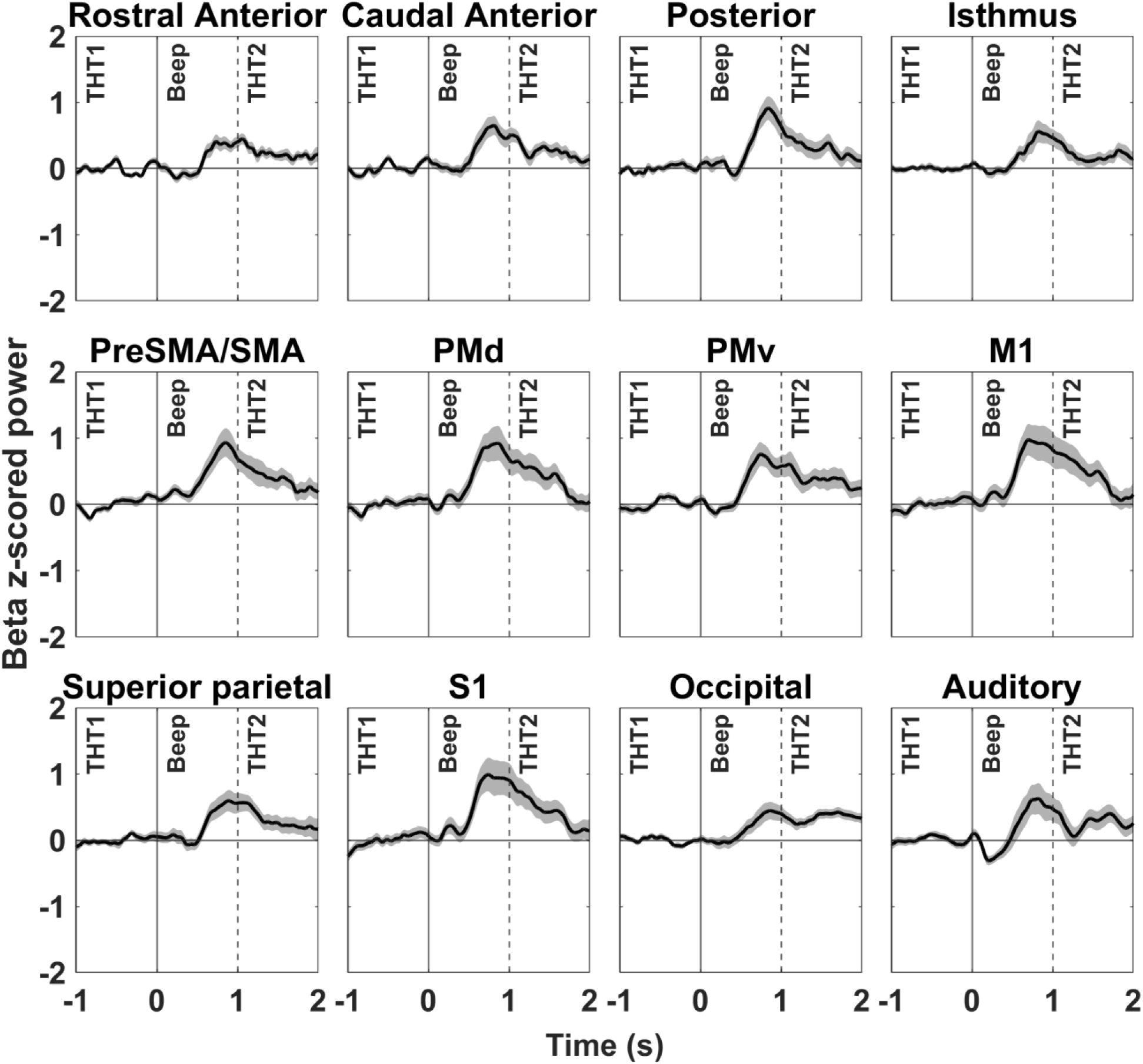
Grand mean (± s.e.m.) time courses of z-score–normalized beta-band (15–29 Hz) power in four cingulate cortex ROIs (top row) and 8 other cortical ROIs (middle and lower rows) in response to the “correct” 1500 Hz Random-Beep auditory tones. Single-trial data are time-locked to Random-Beep tone onset (time 0 s, solid vertical line); tone offset occurred at 1 s (dashed vertical line). Each participant contributed 100 trials, pooled across two Random-Beep blocks. Single-trial data were averaged across the bilateral ROIs for each participant individually and then averaged across participants.

As expected, beta-band MEG power was at baseline before the onset of the Random-Beep tones (Figure 6; time period -1sec to 0sec). The “correct” Random-Beep tones evoked widespread transient increases in beta band power in all 12 ROIs that were largest in M1, S1 and PMd (Figure 6; Supplemental Table 2). The beta response began 400-500ms after tone onset at 0sec, peaked before the offset of the tones at +1sec, and continued to decline towards the pre-tone baseline level during the 1sec THT2 period after tone offset (Figure 6; Supplemental Figure 2).

Alpha-band responses to “correct” Random-Beep tones (Figure 7) were also widespread (Supplemental Table 3, 4), but had more complex and variable temporal profiles than the beta-band responses. The largest response was evoked in the Auditory ROI, that expressed a brisk transient increase in alpha power at the onset of the Random-Beep tone followed by a sharp suppression below baseline that ended just before the offset of the tone at +1sec. A similar but attentuated response was recorded in the PMv ROI, which includes a caudal portion of Broca’s area (c.f., Figure 3). Most other ROIs expressed only a small initial suppression followed by a rebound above baseline near the end of the Random-Beep tone that extended beyond tone offset. The late rebound magnitude was largest in the S1, M1, and Rostral Anterior Cingulate ROIs (Supplemental Table 4).

**Figure 7.**
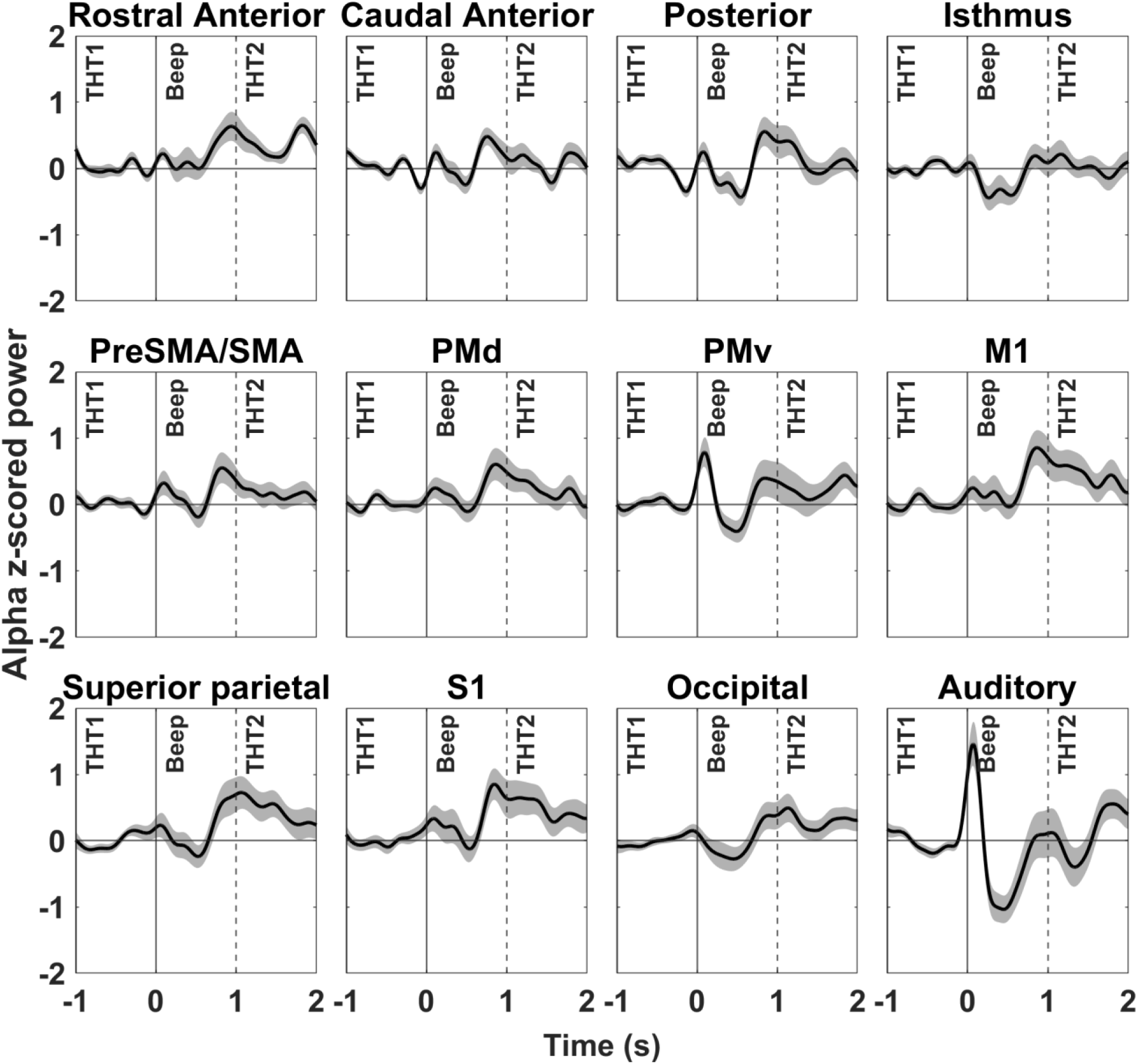
Grand mean (± s.e.m.) time courses of z-score–normalized alpha-band (8− Hz) power in four cingulate cortex ROIs (top row) and 8 other cortical ROIs (middle and lower rows) in response to the “correct” 1500 Hz Random-Beep auditory tones. Same format as Figure 6.

In summary, the “correct” Random-Beep tones evoked spatially organized MEG neural responses in the beta and alpha bands across all ROIs, even though the participants were not actively engaged in the decision-making tasks.

### Post-Feedback-Beep beta and alpha band responses in correct trials across ROIs

We analyzed MEG data from the 12 bilateral pairs of ROIs during the post-decision epochs (THT1, Beep, THT2) of the TFD and CFD tasks (see Methods, Figure 4). The THT1 epoch spanned the 1-second interval from the detection of the motor-report button press that the participants made to indicate their linked perceptual and motor decisions (Figure 4, time –1sec) to the onset of the auditory Feedback-Beep signal (Figure 4, time 0sec). The Beep epoch comprised the subsequent 1-second period from 0sec to +1sec during which participants received a 1500 Hz auditory Feedback-Beep signal indicating a correct decision. The THT2 epoch was the final 1-second delay interval extending from the end of the Beep epoch (+1sec) to the end of the trial (+2sec). Throughout all three epochs, participants continued to press the mouse button that indicated their chosen Target. The main overall finding was that post-Feedback-Beep responses in both bands were much larger in Weak- and Intermediate-evidence trials than in Strong-evidence trials.

### Post-Feedback-Beep Beta Band Rebound

The onset of the 1500Hz correct Feedback-Beep signal (time 0sec, Figure 8) evoked a rapid increase in beta-band power in all 12 ROIs, that began at ∼150-200msec after Feedback-Beep onset and peaked at ∼450-500msec post-Beep onset, before declining toward baseline for the remainder of the Beep and subsequent THT2 epoch. These temporal dynamics were more rapid than the responses evoked by the Random-Beep tones (Supplemental Figure 2A, B). Importantly, the magnitude of the Feedback-Beep-evoked beta rebound was strongly modulated by the evidence strength on which the correct decisions were based in most ROIs and varied substantially across ROIs (Figure 8; Supplemental Table 2).

**Figure 8.**
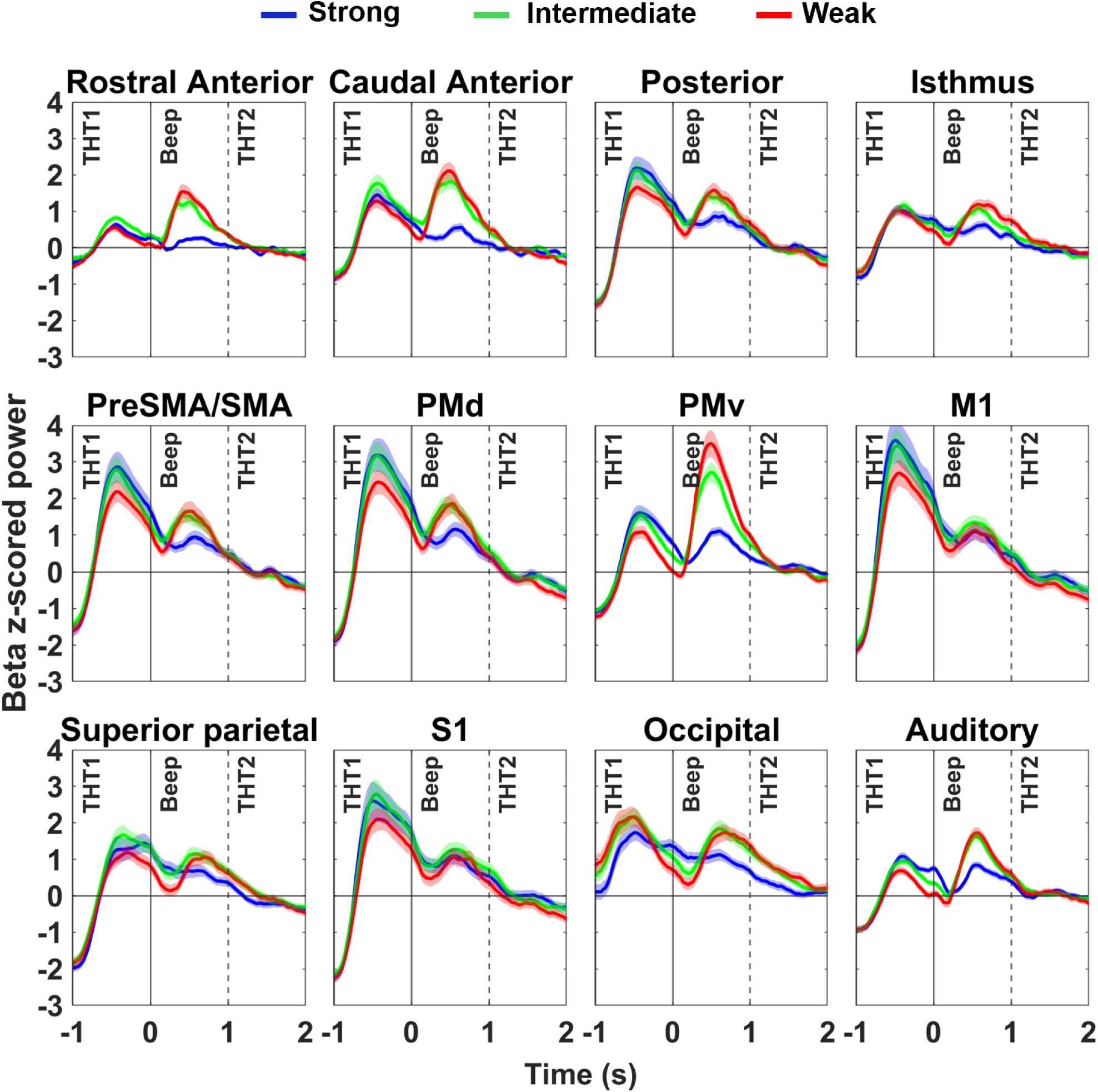
Grand-mean z-score-normalized (±s.e.m.) beta (15–29 Hz) power in four cingulate ROIs (top row) and 8 other cortical ROIs (middle and bottom rows) across the three post-decision epochs (THT1, Beep, THT2), aligned to Feedback-Beep onset (time 0). Only trials with correct decisions were included. Single-trial data were grouped by sensory evidence strength: Strong (blue), Intermediate (green), and Weak (red) (see Methods). Single-trial data were averaged across the bilateral ROIs for each participant individually in each of the TFD and CFD tasks separately and then averaged across both tasks and all participants.

**Table 1.** Permutation test results on differences in post-Feedback-Beep beta-band rebound magnitude in correct trials with different levels of color evidence strength in defined ROIs.

| Permutation test: | p (unadjusted) |  |  | p (FDR-adjusted) |  |  |
| --- | --- | --- | --- | --- | --- | --- |
| Evidence difference: | W-S | I-S | W-I | W-S | I-S | W-I |
| Cingulate ROI: |  |  |  |  |  |  |
| RAC | **0.001** | **0.001** | 0.072 | **0.001** | **0.001** | 0.077 |
| CAC | **0.001** | **0.001** | 0.061 | **0.001** | **0.001** | 0.077 |
| PC | **0.001** | **0.001** | 0.077 | **0.001** | **0.001** | 0.077 |
| IC | **0.001** | **0.002** | **0.002** | **0.001** | **0.002** | **0.008** |
| Cortical ROI: |  |  |  |  |  |  |
| preSMA/SMA | **0.001** | **0.001** | 0.13 | **0.001** | **0.002** | 0.21 |
| PMd | **0.001** | **0.002** | 0.069 | **0.006** | **0.003** | 0.14 |
| PMv | **0.001** | **0.001** | **0.006** | **0.001** | **0.002** | <u>0.050</u> |
| M1 | **0.005** | <u>0.017</u> | 0.576 | **0.006** | <u>0.019</u> | 0.58 |
| SPC | **0.001** | **0.001** | <u>0.029</u> | **0.001** | **0.002** | 0.08 |
| S1 | **0.006** | <u>0.029</u> | 0.258 | **0.006** | <u>0.029</u> | 0.29 |
| OCC | **0.001** | **0.001** | 0.170 | **0.001** | **0.002** | 0.23 |
| AC | **0.001** | **0.001** | <u>0.023</u> | **0.001** | **0.002** | 0.080 |

p (unadjusted): permutation test result for each ROI individually; p(FDR-adjusted): permutation test result for each ROI after False Discovery Rate correction.
p-value format: **p<0.01**; 0.01<p<0.05; p>0.05.
Abbreviations: ROI: region of interest; W-S: Weak – Strong evidence; I-S: Intermediate – Strong evidence; W-I: Weak – Intermediate evidence; RAC: rostral anterior cingulate; CAC: caudal anterior cingulate; PC: posterior cingulate; IC: isthmus cingulate; preSMA/SMA: combined pre-supplemental motor area and supplemental motor area; PMd: dorsal premotor cortex; PMv: ventral premotor cortex; M1: primary motor cortex; SPC: superior parietal cortex; S1: primary somatosensory cortex; OCC: occipital cortex; AC: auditory cortex.

**Table 2.** Permutation test results on differences in post-Feedback-Beep alpha-band suppression magnitude in correct trials with different levels of color evidence strength in defined ROIs. Same format as Table 1.

| Permutation test: | p (unadjusted) |  |  | p (FDR-adjusted) |  |  |
| --- | --- | --- | --- | --- | --- | --- |
| Evidence difference: | W-S | I-S | W-I | W-S | I-S | W-I |
| Cingulate ROI: |  |  |  |  |  |  |
| RAC | <u>0.040</u> | 0.73 | 0.050 | 0.080 | 0.84 | 0.10 |
| CAC | 0.75 | 0.31 | 0.22 | 0.75 | 0.75 | 0.29 |
| PC | 0.70 | 0.38 | 0.61 | 0.75 | 0.75 | 0.61 |
| IC | **0.009** | 0.84 | <u>0.010</u> | <u>0.040</u> | 0.84 | <u>0.05</u> |
| Cortical ROI: |  |  |  |  |  |  |
| preSMA/SMA | <u>0.010</u> | 0.25 | 0.18 | <u>0.03</u> | 0.63 | 0.24 |
| PMd | **0.010** | 0.42 | <u>0.040</u> | <u>0.03</u> | 0.63 | 0.13 |
| PMv | 0.49 | 0.20 | <u>0.050</u> | 0.49 | 0.63 | 0.13 |
| M1 | **0.002** | 0.36 | <u>0.020</u> | <u>0.02</u> | 0.63 | 0.13 |
| SPC | 0.33 | 0.46 | 0.09 | 0.44 | 0.63 | 0.19 |
| S1 | 0.080 | 0.95 | 0.12 | 0.16 | 0.95 | 0.19 |
| OCC | 0.14 | 0.47 | 0.38 | 0.22 | 0.63 | 0.44 |
| AC | 0.44 | 0.72 | 0.58 | 0.49 | 0.83 | 0.58 |

To quantify the effect of color evidence strength on the beta-band responses, we measured the z-scored magnitude of the post-Feedback-Beep beta rebound from the local minimum shortly after tone onset to the subsequent beta rebound peak (see Methods, Figure 4). This measurement was performed separately for the Strong, Intermediate, and Weak color-evidence trial groups, in each of the CFD and TFD tasks for each participant. We then pooled these measurements across tasks and participants.

The Feedback-Beep-evoked beta rebound magnitude decreased as evidence strength increased and associated error rates decreased. It was largest in trials with Weak evidence (Figure 8, red lines; Supplemental Table 2; grand mean 2.433 arbitrary z-score units across all 12 ROIs), smaller in trials with Intermediate evidence (Figure 8, green lines; Supplemental Table 2; grand mean 1.950 z-score units) and was smallest in trials with Strong color evidence (Figure 8, blue lines; Supplemental Table 2; grand mean 1.176 z-score units). Pooled across all 12 ROIs, the beta rebound magnitudes were highly significantly different (p < 0.001) across all three different levels of evidence strength (Supplemental Table 7; one-sample t-test, *ttest*, MATLAB R2023b).

The range of z-scored beta rebound magnitudes across ROIs (smallest to largest) also increased as evidence strength decreased, from 0.98 z-score units in Strong-evidence trials, to 2.04 units in Intermediate-strength trials and 3.08 units in Weak-evidence trials, compared to only 0.9 units in Random-Beep trials (Supplemental Table 2). The increase from Strong-***to Weak-evidence trials was largest in PMv and in the Caudal and Rostral Anterior Cingulate ROIs, and even strong in the Occipital and Auditory ROIs (Figure 8; Supplemental Table 2). Strikingly, the Feedback-Beep beta-band responses in M1 and S1 were among the smallest and least modulated by evidence strength of all cortical ROIs.

For more quantitative comparisons, we conducted permutation tests to compare the distributions of single-participant beta rebound magnitudes between trial groups with different levels of color evidence strength in each ROI individually (Table 1; *permutest* for dependent samples, MATLAB R2023b).

The mean magnitude of the Feedback-Beep-evoked beta rebound was highly significantly larger in every ROI in Weak-evidence trials compared to Strong-evidence trials, including the small difference in M1 and S1 (p << 0.01, unadjusted, Table 1). The mean magnitude was also significantly larger in Intermediate-evidence trials than Strong-evidence trials in all ROIs except M1 and S1, which were significant at only the p = 0.017−0.03 level (unadjusted, Table 1). In contrast, the difference in beta rebound magnitude in Weak-***vs. Intermediate-evidence trials was modestly significant or not significant in most ROIs, except for the Isthmus Cingulate and PMv (Table 2). These results based on unadjusted p-values in each ROI separately largely survived stringent FDR adjustment for multiple comparisons (Table 1).

As already noted, the rank-ordering of beta rebound magnitudes paralleled the increases in objective error rates across trial conditions and was inversely related to evidence strength. Similarly, the differences in beta rebound magnitudes (Supplemental Tables 2, 7) were inversely related to the differences in mean net color evidence in different trials. It was largest (1.26 z-score units) between trials with Weak versus Strong CKBs (44.5 fewer correct-color squares in Weak CKBs), next largest (0.78 z-score units) between Intermediate and Strong CKBs (35.5 fewer squares) and smallest (0.48 z-score units) between Weak and Intermediate CKBs (9 fewer squares).

In contrast, the differences in rebound magnitude did not parallel the differences in error rates between trials with different evidence strength. The largest difference in grand mean beta rebound magnitude occurred between Weak- and Strong-evidence trials that had the largest difference in error rates (34.11%). However, the next largest difference in grand mean beta rebound magnitude occurred between Intermediate- and Strong-evidence trials that had only a 9.33% difference in error rates, whereas the smallest difference in grand mean beta magnitude occurred between Weak- and Intermediate-strength evidence trials that had a 24.78% difference in error rates.

This indicated that the difference in magnitude of the Feedback-Beep-evoked beta rebound responses across trial conditions was rank ordered according to the difference in color evidence on which the participants based their decisions rather than to the differences in their objectively measured error rates for the different CKBs.

### Post-Feedback-Beep alpha band suppression and rebound

In the alpha band, the Feedback-Beep-evoked response was characterized by a transient power suppression shortly after tone onset, followed by a later rebound that peaked before or near the end of the Beep epoch (Figure 4B; Figure 9). Exceptionally, the Auditory ROI exhibited a pronounced transient increase in alpha power at Feedback-Beep onset, similar to its response to the Random-Beep tones (Figure 7). As with the beta band, alpha suppression and rebound appeared stronger in trials with Weak and Intermediate evidence than in Strong evidence trials in several ROIs. Furthermore, similar to the beta band, the onset of the post-Feedback-Beep alpha rebound was earlier and more rapid than the corresponding component of the Random-Beep responses (Supplemental Figure 2C, D).

**Figure 9.**
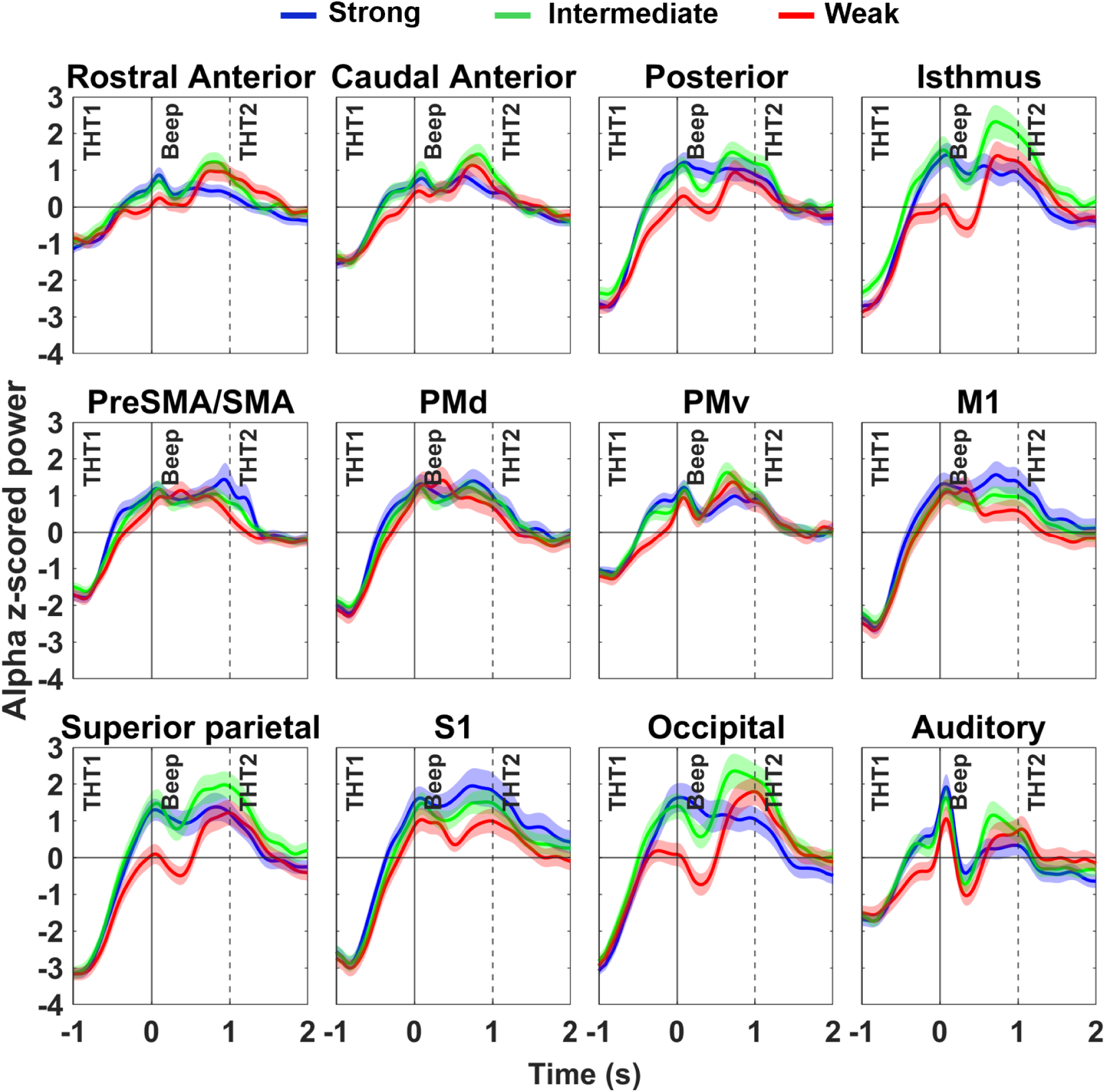
Grand-mean z-score-normalized (±s.e.m.) alpha (8-12 Hz) power in four cingulate ROIs (top row) and 8 other cortical ROIs (middle and bottom rows) across the three post-decision epochs (THT1, Beep, THT2), aligned to Feedback-Beep onset (time 0). Only trials with correct decisions were included. Same format as Figure 8.

Despite the clear suppression visible in the grand mean profiles in Intermediate- and Weak-evidence trials in several ROIs (Figure 9; Supplemental Table 3), there were no significant differences in the measured single-participant alpha suppression magnitudes between different evidence levels when pooled across all 12 ROIs (Supplemental Table 7). Furthermore, when tested individually, only three ROIs showed highly significant differences (p << 0.01; unadjusted) for the Weak–Strong comparison (Table 2), but none maintained that significance after stringent FDR adjustment for multiple comparisons. These modest evidence-dependent effects likely reflect in part the greater difficulty of reliably identifying the initial rebound peak and subsequent suppression trough in the more variable averaged alpha-band profiles of individual trial subsets in individual participants (see Methods).

In contrast, the magnitude of the later Feedback-Beep-evoked alpha rebound was highly significantly larger across all ROIs in Weak-evidence (grand mean 2.521 z-score units) and Intermediate-evidence trials (2.313 z-score units), compared to Strong-Evidence trials (1.757 z-score units), and modestly signficantly larger (p = 0.011) in Weak-than Intermediate-evidence trials (Supplemental Table 7). As was the case for the beta band, the difference in grand mean post-Feedback-Beep alpha band rebound (Weak – Strong: 0.764 units; Intermediate – Strong: 0.557 units; Weak – Intermediate: 0.208 units) was rank ordered according to the difference in color evidence strength in diffferent trials rather than to the differences in experienced objective error rates (Supplemental Tables 4, 7).

The Feedback-Beep-evoked alpha rebound was also significantly stronger in each of the four cingulate ROIs, and the Occipital and Auditory ROIs for both the Weak–Strong and Intermediate–Strong comparisons, which all survived stringent FDR adjustment (Figure 9; Table 3; Supplemental Table 4). The Superior Parietal ROI showed a more modestly significant difference in the Weak–Strong comparison (unadjusted p = 0.005, FDR-adjusted p = 0.01). In contrast, the other sensorimotor ROIs did not show significant differences in post-Feedback-Beep alpha rebound across evidence levels (Figure 9; Table 3; Supplemental Table 4).

**Table 3.**
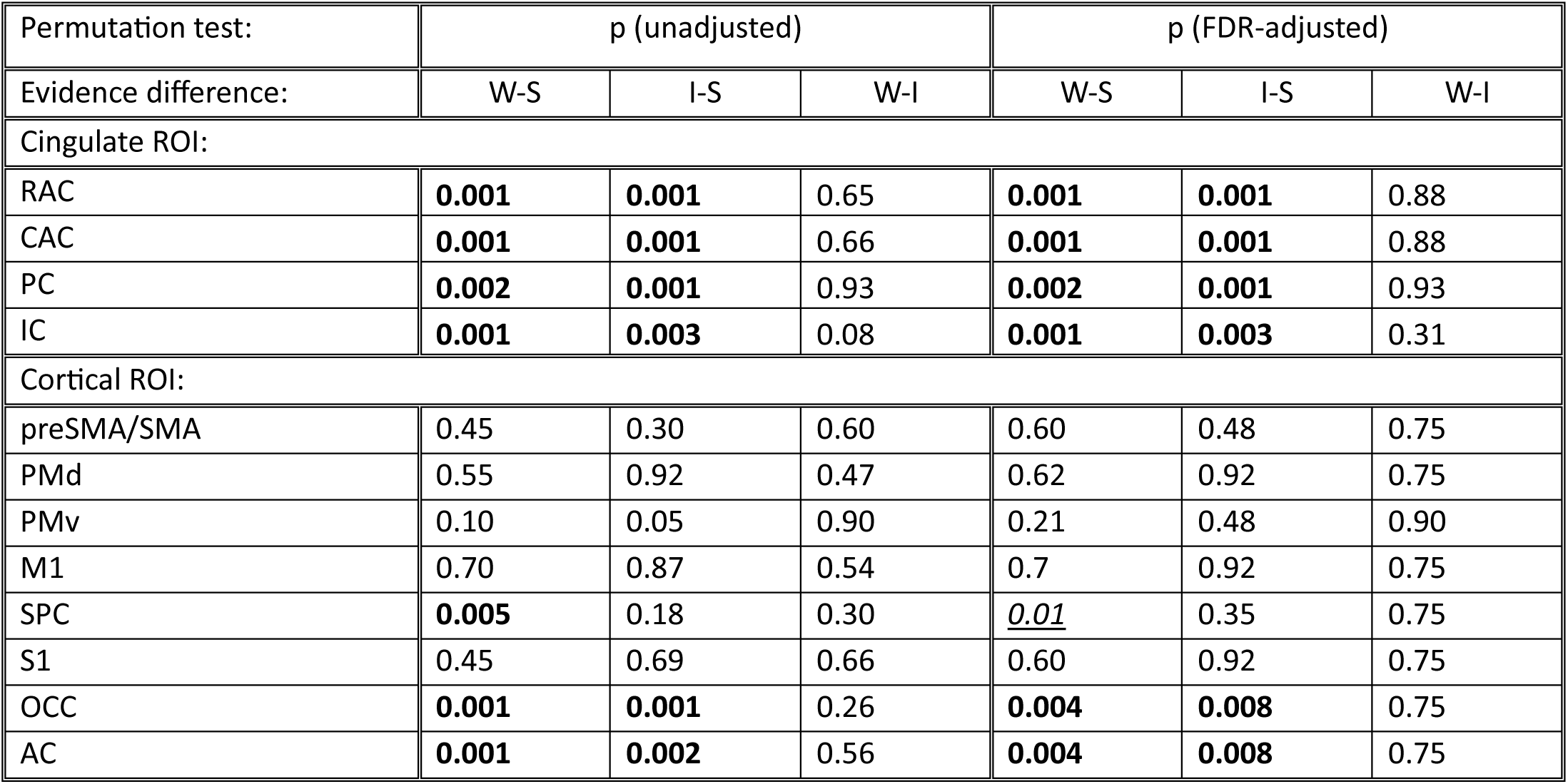
Permutation test results on differences in post-Feedback-Beep alpha-band rebound magnitude in correct trials with different levels of color evidence strength in defined ROIs. Same format as Table 1.

In summary, the alpha band responses following the Feedback-Beep were generally more complex than the beta-band responses and were less modulated by evidence strength, but showed similar trends in effects. Unlike the beta band responses, the evidence-dependent alpha band response modulations were more prominent in Posterior and Isthmus Cingulate and the Occipital and Auditory ROIs than in Anterior Cingulate and premotor ROIs.

### Pre-Feedback-Beep beta and alpha band ROI responses in correct trials

The previous analyses focused on the beta and alpha band responses evoked by the delivery of the Feedback-Beep signals. Another period of interest is the immediately preceding THT1 epoch, which spans the interval between the participants’ button-press report movement and the onset of the Feedback-Beep signal. The multi-colored CKB disappeared at the start of the THT1 epoch, and a small, darker-colored square appeared at the center of the selected Target. Participants continued to press their chosen report button during the THT1 epoch while waiting for the Feedback-Beep signal that would provide them with knowledge of the outcome of their decision. Overall, opposite to the effect of color evidence strength on post-Feedback-Beep responses, the main effect of evidence during the pre-Feedback-Beep THT1 epoch was a strong decrease in alpha band response magnitudes in Weak-evidence trials relative to other evidence levels. The beta band expressed a similar but smaller suppression in Weak-evidence trials.

### Pre-Feedback-Beep THT1 Beta Band Rebound

As already noted, (Figure 4A), beta band power displayed the well-known movement-related suppression below baseline before button press, followed by a rapid power rebound well above baseline during the THT1 epoch that peaked at ∼-600msec to -500msec. It then declined rapidly before the onset of the Feedback-Beep signals at time 0sec in all ROIs (Figure 8). The main exception was the absence of a beta suppression below baseline at the onset of the THT1 epoch in the Occipital ROI. Overall, the post-report-movement rebound was strongest in the M1, PMd, S1, PreSMA/SMA and Posterior Cingulate ROIs, and smallest in the Rostral Anterior Cingulate, Isthmus Cingulate and Auditory ROIs (Figure 8; Supplemental Table 5).

The THT1 beta rebound responses recorded during the -750ms to 0ms window prior to Feedback-Beep onset (Figure 4A) were modestly but systematically smaller in all ROIs in trials with Weak evidence (Figure 8, red lines) compared to Intermediate (green lines) and Strong (blue lines) evidence (Supplemental Table 5). The grand mean of measured THT1 beta rebound magnitudes across all 12 ROIs were highly significantly smaller in Weak-evidence trials (grand mean 1.065 z-score units) than in Strong-evidence (grand mean 1.403 z-score units) and Intermediate-evidence trials (1.396 z-score units), but not between Strong- and Intermediate-evidence trials (Supplemental Table 8). This also differed from the post-Feedback-Beep beta and alpha rebounds, for which the smallest difference was between Weak-evidence and Intermediate-evidence trials (Supplemental Table 7).

Also unlike the post-Feedback-Beep responses, the differences in THT1 pre-Feedback-Beep beta rebounds between trials with different evidence strength (Weak-Strong: -0.338 z-score units; Weak-Intermediate: -0.331 z-score units; Intermediate-Strong: -0.007 z-score units) decreased systematically with the difference in experienced error rates in the different trials (34.11%, 24.78%, 9.33%, respectively), rather than with the difference in net color evidence supporting the correct decision (44.5 squares, 9 squares, 35.5 squares, respectively).

However, when each ROI was tested individually, only PMv (unadjusted p = 0.015) and Auditory (unadjusted p = 0.017) ROIs exhibited a modestly significant difference in THT1 beta magnitude between Weak and Strong evidence trials, whereas the Rostral Anterior Cingulate ROI showed similarly modest differences between Intermediate and Strong evidence trials and between Intermediate and Weak evidence trials (Table 4; permutation test for dependent samples). None of those effects survived stringent FDR correction for multiple comparisons (Table 4).

**Table 4.**
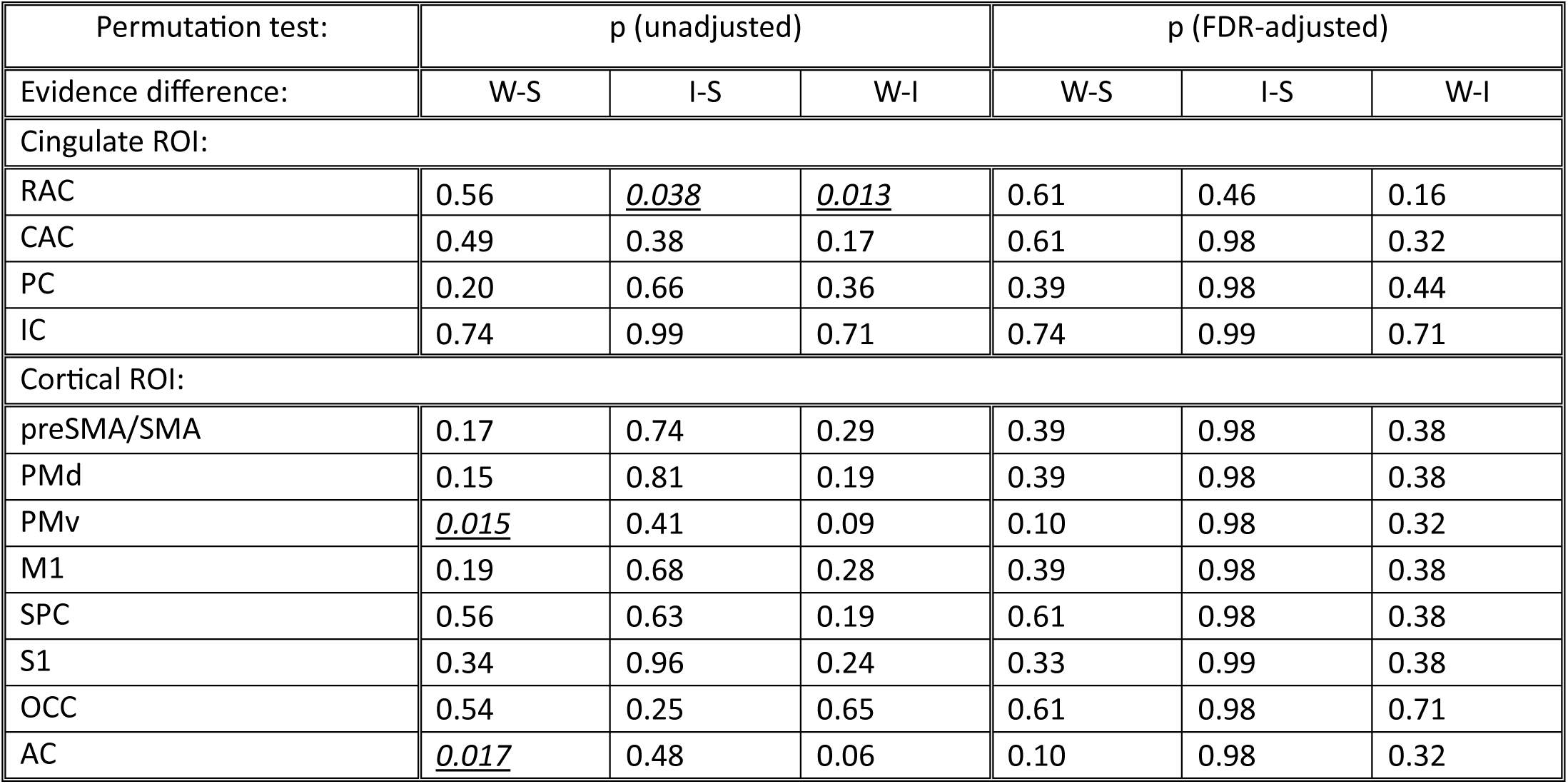
Permutation test results on differences in pre-Feedback-Beep THT1 beta-band rebound magnitude in correct trials with different levels of color evidence strength in defined ROIs. Same format as Table 1.

In summary, there was a highly significant reduction in THT1 beta rebound magnitude between the button-press report movement and the presentation of the Feedback-Beep signal in Weak-evidence trials relative to Strong- and Intermediate-evidence trials when tested across all ROIs, but this effect was largely non-significant in individual ROIs.

### Pre-Feedback-Beep THT1 Alpha Band Rebound

Alpha band activity during the THT1 epoch exhibited an initial movement-related suppression below baseline that typically peaked shortly after the detection of the button press, followed by a rebound that peaked at or shortly after Feedback-Beep onset in most ROIs (Figure 4B, 9). Due to its slower temporal profile than the beta THT1 rebound, we measured average alpha band power in each ROI during a window from −500ms before to +250ms after Beep onset.

The THT1 alpha rebound magnitude was systematically much smaller across all ROIs in trials with Weak evidence (Figure 9, red lines; grand mean 0.170 z-score units) compared to Intermediate (green lines; grand mean 1.314 z-score units) and Strong (blue lines; grand mean 1.518 z-score units) evidence (Supplemental Table 6). These differences were highly significantly different between Weak- and Strong-evidence trials and between Weak- and Intermediate-evidence trials but only marginally significant (p = 0.05) between Intermediate- and Strong-evidence trials (Supplemental Table 8).

Like the THT1 beta rebound responses, the decrease in THT1 pre-Feedback alpha rebound magnitudes paralleled the decrease in color evidence in the CKBs and was inversely related to the associated increases in objective error rates. Also like the THT1 beta band responses, the differences in THT1 alpha rebound magnitude (Weak-Strong: -1.348 z-score units; Weak-Intermediate: -1.144 z-score units; Intermediate-Strong: -0.204 z-score units) decreased systematically with decreases in the experienced error rates in different trials rather than with the difference in net color evidence strength.

In individual ROIs, the rebound was highly significantly smaller in the Posterior Cingulate ROI for Weak-versus Strong-evidence trials (unadjusted p = 0.005; Table 5) and in the Isthmus Cingulate ROI for Weak-***versus Intermediate-evidence trials (unadjusted p = 0.003). Significant differences were also observed in the Superior Parietal ROI for both Weak-***versus Strong-evidence (unadjusted p = 0.008) and Weak-***versus Intermediate-evidence comparisons (unadjusted p = 0.006), and in the Occipital ROI between Weak- and Intermediate-evidence trials (unadjusted p = 0.007). Several other ROIs showed more modestly significant differences (0.01 < unadjusted p < 0.05; Table 5). The incidence and strength of significant effects decreased after stringent FDR correction (Table 5).

**Table 5.**
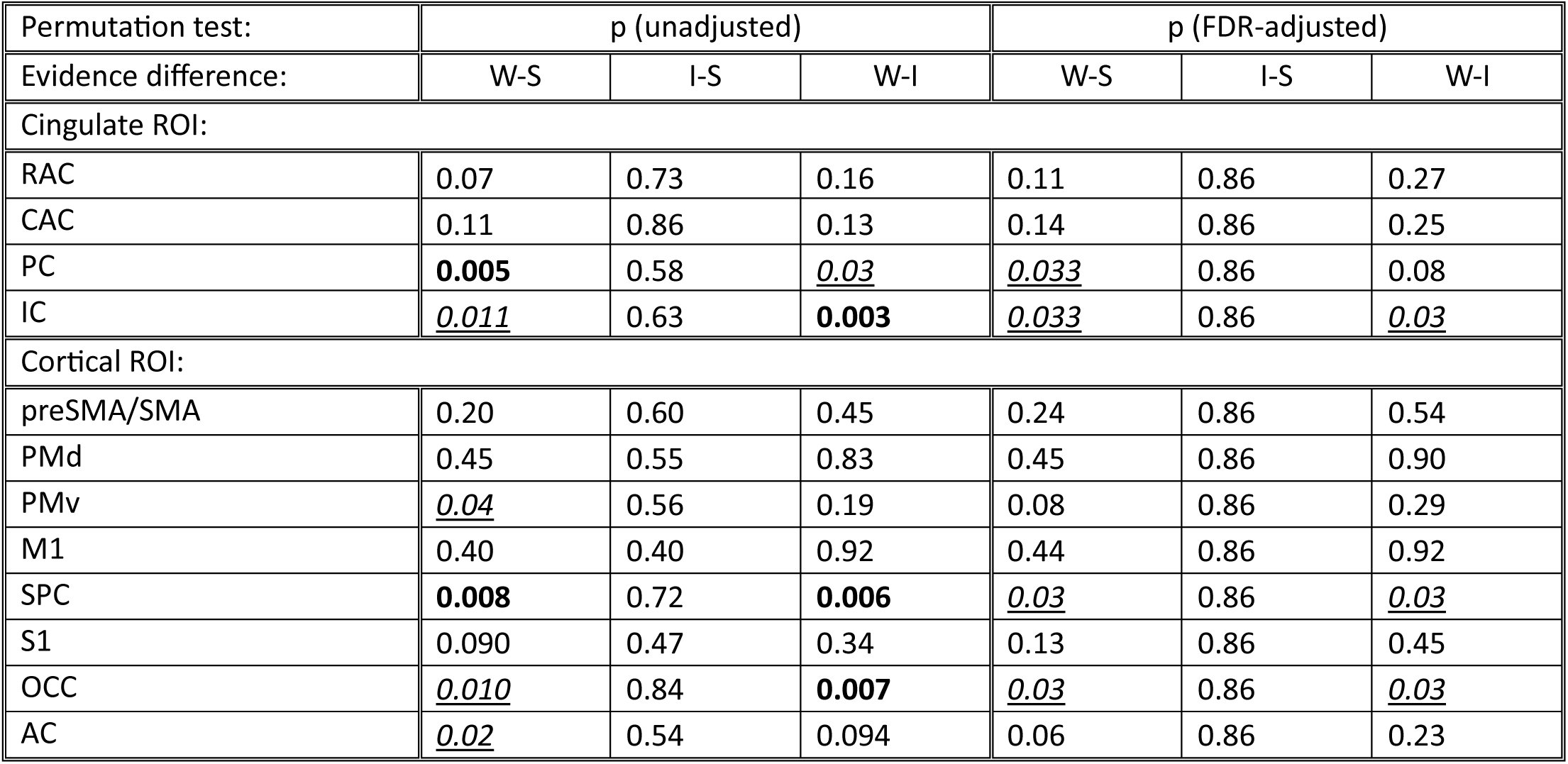
Permutation test results on differences in pre-Feedback-Beep THT1 alpha-band rebound magnitude in correct trials with different levels of color evidence strength in defined ROIs. Same format as Table 1.

Nonetheless, these results indicate that the THT1 alpha band rebound was highly significantly smaller in Weak evidence trials when pooled across all 12 ROIs, and that several ROIs exhibited significantly smaller alpha band rebound magnitudes during the THT1 epoch in correct trials with Weak sensory evidence, while participants awaited the Feedback-Beep signal indicating the outcome of their decision. As was the case for the post-Feedback-Beep alpha band responses, the pre-Feedback suppression of alpha-band rebound was largest in the Posterior and Isthmus Cingulate and Occipital ROIs, and small or absent in motor and premotor cortical ROIs.

### Potential Time Trends within Trial Blocks

The participants performed each task in 4 separate consecutive trial blocks comprising 48 correct trials and variable numbers of incorrect trials. To assure 48 correct trials in each block, we adopted a strategy of re-inserting incorrectly-performed trials later in the pseudo-random trial sequence and repeated them until all trial conditions were performed correctly (see Methods). A consequence of this strategy is that trials with Weak evidence, which had the highest error rate, became increasingly frequent as time progressed within each trial block. This introduced a potential confound about the origin of the evidence-dependent post-decision response modulations. The data analyses up to this point were performed after all the trials in each block were triaged according to the evidence strength in each trial. This segregation implied that the modulations were coupled primarily to processes associated with the CKB evidence experienced in each trial. However, it is also possible that the modulations depended on processes that evolved across time as the relative frequency of Weak-evidence trials increased within each trial block. To assess the effect of this potential confound, we re-analyzed the data after dividing the trials in each block into three sequential sub-blocks (SB1-SB3) with approximately equal numbers of trials, including both correct and incorrect outcomes.

As expected, the incidence of trials with different evidence strength and the rate of correct versus incorrect trial outcomes evolved progressively across the three trial sub-blocks. The frequency of Strong-evidence trials decreased from 32.7% in SB1 to 14.9% in SB3, while the frequency of Weak-evidence trials increased from 33.4% to 60.4% from SB1 to SB3. In parallel, the overall frequency of correct trial outcomes decreased from 85.1% to 74.4% from SB1 to SB3, while the frequency of incorrect outcomes increased reciprocally from 14.9% to 25.6%. Importantly, however, the rate of correct responses in Weak-evidence trials was 64.0% in SB1, 63.0% in SB2 and 62.8% in SB3. This showed that the participants maintained essentially the same success rate in Weak-evidence trials throughout the duration of the trial blocks even though Weak-evidence trials were presented nearly twice as frequently in SB3 than in SB1. They showed no sign of beginning to “guess” when the Weak-evidence trials became increasingly frequent as time progressed in each trial block.

We then tested the stability of the beta and alpha band post-Feedback-Beep rebound responses in correct trials pooled across all 12 ROIs in each of the three trial sub-blocks. Despite the smaller numbers of trials, the response magnitudes were generally similar across trial sub-blocks. For the beta band, rebound magnitudes pooled across all ROIs were statistically identical in SB1, SB2 and SB3 in both Weak-evidence and Intermediate-evidence trials (Supplemental Table 9). In Strong-evidence trials, the beta rebound magnitudes were modestly smaller (p = 0.013) in SB2 compared to SB1, but were non-significantly different between SB1 and SB3, indicating no consistent trend across sub-blocks. For the alpha band, there were no significant differences across sub-blocks in Intermediate-evidence trials. The rebound magnitudes were highly significantly smaller in Strong-evidence trials in SB2 than in SB1, but highly significantly larger in SB3 than SB1. Finally, there was a modest increase in alpha rebound magnitudes (p = 0.014) in Weak-evidence trials in SB2 compared to SB1, but no difference between SB1 and SB3 (Supplemental Table 9). These results also did not indicate a systematic trend across sub-blocks for the alpha band responses.

We also repeated the permutation tests for significant evidence-dependent differences in individual ROIs for each of the three trial sub-blocks. The overall trends in outcomes were consistent with the results for the entire trial blocks (Tables 1, 3), but once again, the incidence of statistically significant outcomes was reduced because of the decrease in single-trial sample sizes (data not shown).

In summary, these findings revealed no compelling evidence for a significant systematic change in the magnitude of post-Feedback-Beep beta and alpha rebound responses across time within each trial block or as the relative frequency of Weak-evidence trials and associated error rates increased within each trial block. These results provide strong support for the conclusion that the evidence-dependent modulations in post-decision responses were primarily coupled to the CKB evidence presented in each trial, independent of its location in the trial block.

### Other Frequency Bands

MEG activity was also examined in the delta (2-4Hz), theta (5-7Hz) and low gamma (30-59Hz) frequency bands. However, they showed relatively minor evidence-dependent modulations of responses evoked by correct Feedback-Beep signals. As an example, responses in the delta band are shown in Supplemental Figures 3 and 4 and Supplemental Table 10. As a result, we did not analyze those data in detail for this study.

## Discussion

We used MEG recordings to investigate the neural dynamics of post-decision outcome-feedback processing in several a priori defined ROIs during a deterministic sensorimotor decision-making task. Participants performed a color-based sensory discrimination task that also required a differential bimanual motor-response decision to report their perceptual decision. Auditory feedback signals presented 1sec after the decision-report movements reliably and accurately indicated whether each decision was correct or incorrect. Our analyses in this report focused on trials with correct feedback signals. Neural responses in incorrect trials and pre-decision activity will be described in future work.

Our primary goal was to examine how the strength of sensory evidence supporting correct sensorimotor decisions modulated neural responses across different MEG oscillatory frequency bands during the post-decision outcome-feedback period. To this end, we contrasted MEG activity in the beta and alpha bands before and after correct Feedback-Beep signals in trials with different levels of color evidence and with responses to Random-Beep tones.

The ROIs were selected in part to test the concept of embodied decision-making. Specifically, in this study, we tested whether nominally “sensorimotor” cortex (M1, S1, PMd, PMv, preSMA/SMA, and Superior Parietal ROIs) would express activity that implicated them in the monitoring of the outcomes of the motor-report actions that they had planned and executed based on different levels of sensory evidence that supported the correct sensorimotor decision.

As stated earlier, the Random-Beep condition cannot be regarded as a pure “sensory” baseline, but it further clarifies the nature of the post-Feedback responses observed during task performance. Although the “correct” Random-Beep tones were physically identical to the correct Feedback-Beep signals, they were presented in the absence of a decision context and therefore lacked evidence-specific decision uncertainty and outcome expectancy. The fact that feedback-locked beta- and alpha-band responses during the task differed systematically from Random-Beep responses indicates that the observed effects cannot be explained by auditory stimulation or learned tone valence alone. Instead, these differences highlight the critical role of decision context and internally generated outcome expectations in shaping cortical responses to outcome feedback.

### Principal Findings

#### Post-Feedback neural activity

A key novel finding of this study was that the presentation of the correct Feedback-Beep tone after a correct decision in the deterministic TFD and CFD tasks evoked a rapid transient +RPE-like increase in MEG power in the beta and alpha bands. Critically, the beta band rebound magnitude was inversely related to the strength of evidence on which the correct decisions were made and the associated expectation of a correct outcome. It was smallest when evidence strength was Strong and participants made almost no incorrect decisions. In contrast, it was largest when color evidence was Weak, participants made the most errors and the expectation of a correct decision was lowest. These evidence-dependent modifications of the post-Feedback-Beep beta band rebound magnitude were most prominent in the more anterior Cingulate ROIs, several premotor ROIs (PMv, PMd, preSMA/SMA, and Superior Parietal), and in the Occipital and Auditory ROIs, but were weak in the M1 and S1 ROIs. Post-Feedback-Beep alpha band rebounds also showed modifications of magnitude that were inversely related to color evidence strength and expectations of correct decisions, but were generally smaller in magnitude, were most prominent in more posterior Cingulate, Superior Parietal, Occipital and Auditory ROIs and spared M1 and the other premotor ROIs.

Furthermore, the rank order of the differences in post-Feedback beta and alpha band response magnitudes across trial conditions was more closely related to the differences in evidence strength between trials than to the differences in experienced error rates. This indicated that the differences in post-Feedback response magnitudes were primarily conditioned by the differences in net evidence that supported the correct decisions between trials and were less modulated by the differences in objective outcome probabilities (error rates) associated with the different levels of evidence. However, because we did not ask participants to report their decision uncertainty on a trial-to-trial basis, we cannot determine whether the rank-ordering of differences in post-Feedback rebound magnitudes might have been better related to their subjective outcome expectations rather than to the objectively measured error rates (Hajcak et al 2005, 2007; Ichikawa et al 2010; Hoy et al 2021).

These Feedback-evoked MEG beta- and alpha-band responses displayed properties expected of positive Reward Prediction Error-like (+RPE-like) signals, defined operationally as correct-outcome responses whose magnitude increased as the expectation of a correct outcome decreased with decreases in evidence strength in a deterministic task context. This increased the magnitude of the prediction error resulting from the same correct outcome in trials with progressively weaker evidence strength.

To our knowledge, these results constitute the first demonstration of +RPE-like MEG signals in sensorimotor cortical regions in human sensorimotor decision-making tasks where outcomes were fully determined by the participants’ evidence-based behavior, rather than probabilistically manipulated to introduce artificial outcome uncertainty or unexpected trial outcomes, as is common in performance-monitoring and RPE paradigms.

Importantly, the present findings support an embodied decision-making framework in a distributed and hierarchical sense, rather than implying that motor cortex per se functions as a reward or reinforcement-learning system. The strongest evidence-dependent post-Feedback response modifications were observed in the beta band in premotor (PMv, PMd, preSMA/SMA) and Superior Parietal ROIs, as well as in anterior Cingulate ROIs, whereas primary motor (M1) and primary somatosensory (S1) cortices exhibited comparatively weaker and less evidence-modulated responses. This revealed that those premotor cortical regions were informed of the successful outcome of the sensorimotor decision to which they contributed, and also the degree to which the positive outcome was expected or not. This pattern suggests a gradient of involvement in outcome evaluation, in which associative sensorimotor regions that integrate perceptual, motor, and contextual information contribute more strongly to post-decision evaluative processes than regions that are more directly involved in movement execution (M1) and movement-evoked somatosensory feedback processing (S1). Thus, embodiment here refers to the engagement of motor-related cortical networks in outcome evaluation, not to a uniform or obligatory role of primary motor cortex in the reward prediction error computation. Indeed, the present results are agnostic as to the specific location of the RPE calculations. The data cannot determine to what degree each ROI may generate the +RPE-like signals or instead expresses a copy of those signals received from neural structures that are more causally responsible for their generation.

The temporal coupling of these MEG events to the Feedback-Beep signals make them an analog of the Feedback-Related Negativity (FRN) and P3/P300 in EEG studies (Miltner et al. 1997; Holroyd and Coles 2002; Holroyd et al. 2003; Hajcak et al. 2005; 2007; Cohen et al. 2007; Marco-Pallarés et al. 2008; HajiHosseini et al. 2012), rather than the Error-Related Negativity (ERN) (Gehring et al. 1993; Holroyd and Coles 2002; Frank et al. 2005). Our finding of +RPE-like responses to correct outcome feedback signals is consistent with prior EEG and MEG studies of FRN or P3/P300 potentials in a wide range of tasks with and without probabilistic reward policies (Holroyd et al. 2003; 2004; 2008; Hajcak et al. 2005; 2007; Cohen and Ranganath 2007; Marco-Pallarés et al. 2008; Philiastides et al. 2010; HajiHosseini et al. 2012; Ferdinand et al. 2012; Bai et al. 2015; Fouragnan et al. 2017; Walentowska et al. 2019; Billeke et al. 2020; Talmi et al. 2012; Thomas et al. 2013).

Most relevant to the current study, Cohen et al. (2007), Marco-Pallarés et al. (2008), HajiHosseini et al. (2012), and HajiHosseini and Holroyd (2015a, b) all found increased beta band power after positive outcome feedback in tasks with probabilistic outcome policies, consistent with the beta band rebounds in correct trials in this study. However, unlike the current findings, some of the studies did not find modulations of the beta response magnitudes by differences in positive outcome probabilities (HajiHosseini and Holroyd 2015a, b). In contrast, HajiHosseini et al. (2012) reported that beta response magnitude in response to positive feedback was large in trials with a low probability of a positive outcome in a gambling/guessing task, but much weaker in trials with a high probability of a positive outcome. This is entirely consistent with our findings of large beta rebound magnitudes in trials with Weak evidence, but small rebounds in trials with Strong evidence, but the different outcome expectations in the HajiHosseini et al. (2012) study were imposed by the probabilistic reward policy.

#### Pre-Feedback neural activity

In contrast to the post-Feedback responses, during the 1sec THT1 delay period between the completion of the decision-report button-press movement and the presentation of the correct Feedback tone, the magnitude of the post-movement alpha rebound response was highly significantly smaller in Weak-evidence trials than in Strong- and Intermediate-evidence trials. This effect was most prominent in the more posterior Cingulate ROIs, and the Occipital and Superior Parietal ROIs, but not in motor or premotor ROIs. A similar effect was expressed in the beta band, but was smaller in magnitude.

Furthermore, the differences in beta and alpha band magnitudes in the pre-Feedback THT1 epoch scaled differently as a function of differences in evidence strength and outcome expectations than in the post-Feedback epochs. The rank order of the differences in pre-Feedback suppression magnitude across trial conditions was more closely related to the differences in experienced error rates between trials with different levels of evidence strength than to the differences in evidence strength themselves. This suggests that differences in outcome expectations had a stronger effect on the pre-Feedback response modifications than evidence strength differences per se.

These pre-Feedback response modifications while the participants were waiting to receive feedback about their decision outcome are not RPE-like signals because the participants had not yet received knowledge of results and so could not calculate a prediction error. Similarly, they should not be interpreted as direct evidence of predictive coding during the THT1 epoch. Rather, we interpret these pre-Feedback alpha- and beta-band effects as correlates of internal decision confidence and outcome expectations that were maintained or elaborated after the decision-report movement while participants awaited outcome feedback (Kepecs et al. 2008; Kiani and Shadlen 2009; Kepecs and Mainen 2012; Drugowitsch et al. 2014; Kiani et al. 2014; van den Berg et al. 2016; Zylberberg et al. 2016; Löffler et al. 2023; Pleskac and Busemeyer 2010; Pouget et al. 2016; Boldt and Yeung 2015; Murphy et al. 2015; Lak et al., 2017; Grogan et al. 2023). These pre-Feedback signals likely reflect a combination of processes, including sustained representations of evidence stength, decision uncertainty and outcome expectations, heightened attentional engagement toward upcoming outcome feedback information (see below), and preparatory neural states for processing feedback, rather than an explicit prediction signal in the sense of formal Reinforcement Learning theory. These effects were expressed most strongly in the alpha band in trials in which participants had made their decision on the basis of the CKBs with the weakest sensory evidence for the correct decision.

Within this framework, sensory evidence strength influences decision confidence, which in turn shapes expectations about decision outcomes. These internally generated expectations are expressed during the post-decision, pre-feedback interval as modulations of alpha- and beta-band activity. Upon receipt of outcome feedback, these expectations are then evaluated against the experienced outcome relative to the evidence on which the decision was based, giving rise to the post-feedback beta- and alpha-band responses that scale with expectancy violation and resemble a +RPE-like signal. This temporal sequence provides a coherent account linking sensory evidence, confidence, pre-feedback neural state, and post-feedback outcome evaluation without requiring that pre-Feedback activity itself encodes a formal prediction error.

### Sensory Evidence, Task Performance, Decision Confidence and Outcome Expectations

Evidence-dependent estimates of decision confidence and outcome expectation are critical components of the proposed RPE computations in these tasks (Kepecs et al. 2008; Kiani and Shadlen 2009; Kepecs and Mainen 2012; Kiani et al. 2014; van den Berg et al. 2016; Zylberberg et al. 2016; Löffler et al. 2023; Pleskac and Busemeyer 2010; Boldt and Yeung 2015; Murphy et al. 2015; Lak et al., 2017; Grogan et al. 2023). The participants in this study appeared to experience these introspective metacognitive states while performing the tasks.

Behavioral performance in the tasks was strongly modulated by the strength of color sensory evidence in the CKBs (Coallier and Kalaska 2014; Coallier et al. 2015; Wang et al. 2019). Participants made virtually no errors in Strong-evidence trials, whereas error rates increased progressively in Intermediate- and Weak-evidence trials (Figure 5). We did not ask participants to report their decision confidence either directly or indirectly on a trial-by-trial basis, as has often been done in previous studies (Kepecs et al. 2008; Kepecs and Mainen 2012; Kiani and Shadlen 2009; Pleskac and Busemeyer 2010; Kiani et al. 2014; Sanders et al. 2016; Zylberberg et al. 2012; 2014; Moran et al. 2015; Boldt and Yeung 2015; van den Berg et al. 2016; Zylberberg et al. 2016; Bang and Fleming 2018; Löffler et al. 2023). Nevertheless, post-experiment debriefing of the participants confirmed that they were aware of this performance trend. Most reported feeling very confident in their decisions during Strong-evidence trials, but reported considerably lower confidence in Weak-evidence trials prior to receiving outcome feedback. Several participants even remarked that they felt as though they were guessing in Weak-evidence trials, despite achieving success rates that were above chance.

These anecdotal self-reports provide strong indirect evidence that participants experienced a lower degree of decision confidence in Weak-evidence trials and had a correspondingly lower expectation of a correct outcome while waiting for the Feedback-Beep signals compared to trials with stronger sensory evidence. That expectation was presumably shaped by each participant’s accumulated history of performance during the training and MEG recording sessions, when experienced outcomes varied systematically with sensory evidence strength in these deterministic tasks (Dehaene et al. 2021; Drugowitsch et al. 2014; Pouget et al. 2016; Philiastides et al. 2010; Bai et al. 2015; Fouragnan et al. 2017).

This inferred link between evidence strength and outcome expectations was further corroborated by the effect of evidence strength on the participants’ RTs in the TFRT task during training (Supplemental Figure 1), which were significantly longer in trials with Weak evidence than Strong evidence (Coallier and Kalaska 2014; Coallier et al. 2015). Many studies have reported a strong correlation between decision duration, decision confidence and reward expectation (Kepecs et al. 2008; Drugowitsch et al. 2014; Kiani and Shadlen 2009; Kiani et al. 2014; van den Berg et al. 2016; Zylberberg et al. 2016; Löffler et al. 2023). Further behavioral support for evidence-dependent changes in decision confidence comes from studies that showed that participants are more likely to change their mind and reverse their decision in trials in which evidence was weak (Resulaj et al. 2009; Coallier and Kalaska 2014; Atiya et al. 2020) or opt out of a trial prematurely rather than waiting to receive feedback about decision outcomes (Kepecs et al. 2008; Kiani and Shadlen 2009).

### Decision confidence, outcome uncertainty, directed attention and enhanced cortical processing

We described the evidence-dependent modulations of the magnitude of pre-Feedback-Beep and post-Feedback-Beep beta and alpha rebounds as neural correlates of processes coupled to post-decision performance monitoring that were influenced by variations in the level of decision confidence and outcome expectations, and that in turn influenced RPE calculations. However, another potential contributing factor could be evidence-dependent trial-to-trial changes in the level of attention that the participants paid to the Feedback-Beep signals, beyond any spontaneous attentional drift across time, because they provided more salient information about task performance as evidence strength and decision confidence decreased (Kepecs and Mainen 2012; Drugowitsch et al. 2014; Pouget et al. 2016). In Strong-evidence trials, the sensorimotor decision was easy and the participants were very confident about the likely success of their decisions, since they experienced only a 0.09% mean error rate in those trials. As a result, they expected to receive a correct Feedback-Beep signal at the end of Strong-evidence trials, and its reception provided very little new information about the trial outcome beyond confirmation that it was correct as expected. It is possible, therefore, that the participants paid a low level of attention in Strong-evidence trials while waiting for the Feedback-Beep signal and while processing it after its presentation. In contrast, in Weak-evidence trials, the sensorimotor decision was more challenging and the participants were less confident about the likely success of their decisions because they experienced a 34.2% mean error rate in those trials. In that situation, the Feedback-Beep signal provided much more salient novel information about the outcome of the trial and the participants may have paid more attention while waiting for the signal and while processing it after it was presented.

This raises the possibility that evidence-dependent trial-to-trial fluctations in the level of attention of the participants to the Feedback-Beep signals could have been the causal agent for the beta and alpha band MEG activity modulations that we observed during the pre-feedback and post-feedback epochs in the tasks (Apitz and Bunzeck 2012; Thomas et al. 2013). Consistent with this hypothesis, suppression of alpha band activity has often been implicated in the regulation of visual attention and cortical excitability (Foxe and Snyder 2011; Jensen and Mazaheri 2010; Haegens et al. 2011b; Klimesch 2012; Van Diepen et al. 2019; Vanni et al. 1997; Kelly et al. 2006; 2009; Gould et al. 2011; Samaha et al. 2017; 2020). This could account for the strong suppression of pre-Feedback alpha rebound power during the THT1 epoch in Weak-evidence trials, which was particularly prominent in the Occipital ROI. However, this would not account for the enhanced post-Feedback alpha rebound magnitude in Weak-evidence trials in the Occipital and Isthmus Cingulate ROIs, which would imply lower attention to the evaluation of the correct Feedback-Beep tones when successful outcomes were the least expected.

We do not propose that evidence-dependent attentional modulations play no role in outcome-feedback processing in these deterministic tasks. It is likely that the different evidence-dependent levels of decision difficulty, decision confidence and expectations of successful outcomes experienced by the participants during the sensorimotor decision-making tasks were in turn the causal agents of any evidence-dependent fluctations in the level of attention across trials. In this scenario, attentional engagement is downstream of internally generated estimates of decision confidence and outcome expectation. Irrespective of their causal relationship, this suggests that introspective evidence-dependent metacognitive inferences associated with the sensorimotor decision-making processes and evidence-dependent fluctuations in the level of attention paid to the information provided by the Feedback-Beep signals were tightly coupled and confounded in these tasks. Changes in task design could help to decouple these different cognitive processes in future studies. Nevertheless, these considerations imply that any potential evidence-dependent fluctations in the level of attention that could have influenced the observed MEG activity are a product of and a surrogate measure for metacognitive inferences about decision confidence, the expected outcomes of the sensorimotor decisions, and Reward Prediction Errors. In this framework, attention is not an independent confound but one mechanism through which expectancy-dependent outcome evaluation is performed, giving rise to the observed +RPE-like post-Feedback beta- and alpha-band responses.

### Deterministic versus probabilistic tasks

The deterministic feedback policy of the sensorimotor decision-making tasks used in this study increases the ecological validity of our findings for many everyday behavioral contexts in which outcome feedback is solely and causally dependent on the accuracy of the decisions. In that context, the decision-maker can reliably associate evidence quality and decision difficulty with the probability of decision outcome expectations, and with the degree to which a positive outcome deviates from the expected outcome as decision difficulty increases. This allows the decision-maker to formulate a reliable estimate of reward prediction errors that they can use to optimize future performance in similar situations, such as maximizing the probability of correct decisions in subsequent trials with Weak evidence strength in our tasks.

However, many tasks that have been used to study RPEs employed probabilistic feedback schedules to create and manipulate artificial discrepancies between evidence, decisions, outcome expectations and experienced outcomes. In such tasks, subjects receive occasional deceptive false negative and false positive outcome feedback in randomly chosen correct and incorrect trials, respectively, dictated by the probabilistic reward policies.

This strategy of using deceptive feedback to generate artificial unexpected outcomes yields misleading reward prediction errors that do not accurately reflect task performance and introduces potentially important extraneous elements of uncertainty into performance monitoring that are related to the degree of veracity of the performance information provided by the feedback signal (Holroyd et al., 2002, 2009; Ferdinand et al 2012). Specifically, was the decision truly correct or incorrect as signaled by the feedback, or did the feedback deceptively indicate the opposite outcome? Alternatively, was the feedback accurate but the subject’s outcome expectation was incorrect, or was the expectation valid but the feedback was unreliable? This raises concerns about the degree to which any observed RPE-like signals in probabilistic tasks reflect the uncertainty and expectations associated with the decision process versus the uncertainty associated with the veracity of the feedback signals. This will muddle the interpretation of any observed RPE-like responses (Holroyd et al., 2009; Ichikawa et al. 2010; Ferdinand et al., 2010). Moreover, this uncertainty reduces the utility of any computed RPE for the decision-maker to improve their task performance because a deceptive feedback signal will yield counterproductive adjustments. This might even discourage participants from attempting to make accurate RPE calculations because they may have little constructive impact on their task performance (Oliviera et al. 2007; Hajcak et al., 2007; Holroyd et al., 2009; Ferdinand et al 2012). This might have also contributed to the diversity of results about RPEs in studies that have used probabilistic reward policies.

This does not mean that tasks with probabilistic reward policies are unlearnable. A task might accord valid positive feedback in 75-80% of correct trials chosen at random, but invalid positive feedback in 20-25% of the incorrect trials chosen at random and still be learnable (Holroyd and Coles, 2002; Hajcak et al., 2005, 2007; Oliviera et al., 2009; Holroyd et al., 2009; Ichikawa et al 2010; Ferdinand et al 2012; HajiHosseini and Holroyd 2015a). A critical factor may be the relative frequency of invalid positive feedback in incorrect trials; if they are never presented, a task can still be learned even when the incidence of valid positive feedback in correct trials is less than 50% (Holroyd and Coles 2002; Holroyd et al 2009; HajiHusseini and Holroyd 2015a). Critically, post-feedback responses have been shown to differ when feedback policies render tasks more or less learnable (Holroyd and Coles, 2002; Hajcak et al., 2005, 2007; Oliviera et al., 2009; Holroyd et al., 2009; Ichikawa et al 2010; Ferdinand et al., 2012), which likely reflect at least in part the uncertainties in performance evaluation introduced by the probabilistic feedback policies.

In contrast, outcome feedback signals that provide information that is a veridical reflection of the participant’s performance are optimal in reinforcement learning and performance monitoring. Deterministic tasks should reveal neural events associated with metacognitive performance monitoring and feedback processing that are dependent completely on the participants’ introspective assessment of their own decision-making. They avoid the added complexity and confounds of other internal neural processes that may be evoked by probabilistic tasks to assess the potentially capricious influence of an external agent (the experimenter) beyond the control of the participants, who metaphorically roles dice to determine what trial “outcome” will be transmitted to the participants despite their best efforts to make correct choices.

Our deterministic task design avoided any confounding neural processes resulting from those added sources of uncertainty in performance monitoring and feedback processing. The degree to which the participants’ expectations of correct or incorrect outcomes were shaped entirely by their individual ability to make correct decisions using the different levels of color sensory evidence in different trials. The observed MEG events reflected neural processes implicated in the processing of veridical feedback about task performance, unencumbered by any extraneous uncertainty about the veracity of the outcome feedback signals.

### Other Potential Confounds

Evidence-dependent post-Feedback beta and alpha rebound magnitude modulations displayed properties consistent with a +RPE-like computation. Importantly, these evidence-dependent modulations were dissociated from the sensory properties of the feedback signals, because all correct decisions were confirmed by the identical 1500Hz auditory tone. Furthermore, the timing and magnitude of Feedback-Beep-evoked responses during the tasks differed significantly from responses to physically identical 1500Hz Random-Beep tones, highlighting their task specificity and relevance to performance evaluation, beyond mere detection and coding of the physical properties of the feedback stimuli, or any automatic recall of the learned valence meaning (“correct”) of the 1500Hz beep tones. These findings about the effects of sensory evidence strength and outcome expectation on post-decision beta and alpha band activity are consistent with prior reports about the effect of response certainty on beta-band activity in the sensorimotor cortex (Donner et al. 2009; Doyle et al. 2005; Grent-’t-Jong et al. 2014; Tzagarakis et al. 2010; 2015; van Helvert et al. 2021).

The participants continued to press down on the decision-report button throughout the post-decision THT1, Beep and THT2 epochs. This rules out any explanation that the evidence-dependent post-decision response modulations were a direct consequence of evidence-dependent differences in overt motor activity or proprioceptive feedback. Instead, these post-decision signals likely reflect centrally generated, evaluative processes related to outcome monitoring.

The post-decision response modulations also cannot be driven directly by the physical differences in the visual sensory evidence because the CKBs that contained the color evidence on which the subjects made their decisions disappeared the moment that the participants made their motor-report button press movement at the end of the Go/RT epoch, that started the THT1 epoch. This suggests that the evidence-dependent modulations of the responses to the Feedback-Beep signals must reflect features of the immediately preceding decision-making process in each trial that continue after the decision-report movement is made or are retained in short-term working memory and recalled when the Feedback-Beep signal is presented. This could include continued processing of the color evidence that is still in the processing pipeline after the CKBs disappeared (Resulaj et al. 2009; Burk et al. 2014), a short-term memory of the strength of evidence on which the participants had just made their decision, along with any associated introspective metacognitive estimates of decision confidence and successful outcome expectation (Kepecs et al. 2008; Kiani and Shadlen 2009; Kepecs and Mainen 2012; Kiani et al. 2014; van den Berg et al. 2016; Zylberberg et al. 2016; Löffler et al. 2023; Pleskac and Busemeyer 2010; Boldt and Yeung 2015; Murphy et al. 2015; Grogan et al. 2023).

### Functional Implications of the Evidence-dependent Post-Decision Response Modulations in Different ROI Groups

The **cingulate cortex** has often been implicated in performance monitoring, decision evaluation, error detection and adaptive control (Botvinick et al. 1999; 2004; Bush et al. 2000; Carter et al. 1998; Holroyd and Coles 2002; Shenhav et al. 2013; Cheyne et al. 2012). Consistent with those roles, our results showed +RPE-like cingulate responses to positive Feedback-Beep signals that confirmed a correct decision, in particular stronger responses that signalled an outcome that was better than expected when a correct decision was based on Weak color evidence. Notably, the Caudal Anterior and Rostral Anterior Cingulate ROIs exhibited the largest beta rebound following correct outcome Feedback-Beep signals in trials with Weak and Intermediate evidence (Supplemental Table 2), consistent with its proposed role in integrating internal confidence with external performance feedback (Carter et al. 1998; Ullsperger et al. 2014).

The alpha rebound following the Feedback-Beep signal also expressed a +RPE-like modulation that was statistically robust across all cingulate ROIs (Table 3) but was strongest in the Posterior and Isthmus Cingulate ROIs (Supplemental Table 4).

These findings support a valence-sensitive, context-dependent role for the cingulate cortex, in which lower decision confidence results in stronger feedback-evoked evaluative responses following a correct outcome. These results align with the hypothesis that the cingulate cortex activity expresses an evidence-dependent RPE or outcome expectancy violations, even though the experienced outcomes in the selected trials in this study were all positive (Fouragnan et al. 2018; Holroyd et al. 2003; Talmi et al. 2012).

Random-Beep tones evoked modest beta and alpha increases in the same cingulate ROIs. This indicated that these ROIs also likely responded to the auditory salience or learned valence of the tones outside of the task context, but that the task-embedded Feedback-Beep signal strongly contextualized and modulated the magnitude of the neural response, consistent with models of adaptive control and performance monitoring (Holroyd and Coles 2002).

Several **cortical sensorimotor ROIs** expressed post-Feedback responses that reflected outcome feedback processing modulated by prior sensory evidence and outcome expectations, that go beyond their traditional roles in movement planning and execution (An et al. 2019; Brasted and Wise 2004; Buch et al. 2006; Derosiere et al. 2025; Kapogiannis et al. 2008; Marshall et al. 2009; Ramakrishnan et al. 2017; Ramkumar et al. 2016; Roesch and Olson 2003). We found strong +RPE-like beta-band responses to the correct Feedback-Beep signals in the PMv, PMd, preSMA/SMA and Superior Parietal ROIs. The PMv ROI exhibited the largest post-Feedback-Beep beta rebound and the strongest evidence-dependent modulation. PMv is a region implicated in higher-order hand control, sensorimotor decision-making, and non-motor cognitive functions, including action observation and mirror neuron activity (Binkofski and Buccino 2006; Chouinard and Paus 2010; 2006; Ramkumar et al. 2016; Rizzolatti et al. 2002; Roesch and Olson 2003; Pardo-Vazquez et al. 2008; Martinez-Garcia et al. 2015; Romo and de Lafuente 2013), making it a likely site for the integration of motor and higher-order cognitive signals. It is also noteworthy that the PMv ROI includes the caudal part of Broca’s area (Chouinard and Paus 2006; 2010), which raises the speculative possibility that part of the strong +RPE-like responses expressed in that ROI reflected sub-vocal processes related to recognition of the learned valence value (“correct”) of the 1500Hz Feedback-Beep signals. The +RPE-like post-Feedback-Beep beta band responses in these sensorimotor ROIs indicate that they contribute to the post-feedback monitoring of task performance, which is one aspect of the larger process of embodied cognition underlying the control of voluntary movements. This implicates them in evaluative processes traditionally associated with higher-order cognitive and limbic structures.

In contrast to the beta band responses, alpha band rebounds after the Feedback-Beep were not significantly modulated by sensory evidence strength in the sensorimotor ROIs, except the Superior Parietal cortex. The absence of a strong alpha rebound modulation in the other sensorimotor ROIs indicates that the neural activity underlying the +RPE-like signals in the cortical sensorimotor network was more prominently captured in the beta band.

The relatively weak +RPE-like post-Feedback-Beep beta and alpha rebound modulations in the M1 ROI suggests that it makes a relatively limited contribution to the post-feedback monitoring of task performance.

The **primary occipital and auditory sensory** ROIs expressed +RPE-like post-Feedback beta and alpha band responses that may reflect modulation of the activation state of the sensory cortical circuits by internal signals concerning directed attention, outcome expectations and performance monitoring, beyond their traditional roles in passive processing of sensory input (FitzGerald et al. 2013; Fritz et al. 2007; Vanni et al. 1997; Kelly et al. 2006; 2009; Foxe and Snyder 2011; Gould et al. 2011; Haegens et al. 2011b; Klimesch 2012; Van Diepen et al. 2019; Samaha et al. 2017; 2020). Our findings provide further evidence that primary sensory cortices are not merely passive processors of afferent sensory inputs, but also exhibit context-dependent oscillatory dynamics modulated by outcome probabilities, reward expectations and outcome feedback (Pessoa and Engelmann 2010; Bunzeck et al. 2011; Apitz and Bunzeck 2012; 2014; Thomas et al. 2013; Bach et al. 2017; Bastos et al. 2015; Meindertsma et al. 2017).

This is further supported by the finding that they also expressed an evidence-dependent suppression of the pre-Feedback alpha band rebound, which could be a possible index of decision confidence or expected outcome before the reception of the Feedback-Beep signals. Consistent with that interpretation, the rank order of differences in the degree of suppression of the pre-Feedback THT1 alpha rebound were closely related to differences in experienced error rates across trial conditions. The lower confidence and heightened expectation of an incorrect outcome could also result in enhanced attentive engagement while awaiting the Feedback-Beep signals in Weak evidence trials, which could also account for the alpha band suppression (Foxe and Snyder 2011; Jensen and Mazaheri 2010; Klimesch 2012; Van Diepen et al. 2019; Vanni et al. 1997; Kelly et al. 2006; 2009; Gould et al. 2011; Samaha et al. 2017; 2020).

The occipital cortex is primarily involved in the initial processing of the color-based visual stimuli (CKBs) which determined the correct decision in each trial. The auditory cortex, by contrast, represents the acoustic features of the Feedback-Beep signals which convey outcome information at the end of each trial. Nevertheless, both ROIs exhibited evidence-dependent pre- and post-Feedback response modulations. Interestingly, the Occipital ROI expressed those effects even though the CKBs that had presented the color evidence had disappeared when the participants made their motor-report movements 1sec before the onset of the Feedback-Beep signals. This suggests that these task-critical sensory areas are informed about pre-Feedback expectations and post-Feedback outcomes of the sensorimotor decision process to which they contributed, possibly through top-down signals reflecting internal performance monitoring. Potential sources of those signals include the Posterior Cingulate and Isthmus Cingulate ROIs that exhibited significantly diminished pre-Feedback-Beep alpha rebounds in Weak-evidence trials, and the Caudal Anterior and Rostral Anterior Cingulate ROIs that exhibited large post-Feedback-Beep beta rebounds in correct trials with Weak and Intermediate evidence.

These findings are consistent with a reentrant model of outcome evaluation, in which evaluative cortical hubs such as the cingulate cortex send modulatory signals to task-relevant primary sensory areas, updating their activation state based on the context-dependent interpretation of outcome feedback. Such a model supports the inclusion of early sensory regions in cognitive performance monitoring circuits, especially under conditions of low perceptual or decision-related confidence in deterministic decision-making tasks (Bastos et al. 2015; Meindertsma et al. 2017).

In contrast, the **S1** ROI displayed the weakest beta and alpha rebound responses of the three primary sensory ROIs and the smallest modulation by sensory evidence strength. This indicated that S1 had a limited role in metacognitive processes related to evidence-dependent outcome estimation, internal confidence and outcome evaluation in the TFD and CFD tasks (Molenberghs et al., 2016; Vaccaro and Fleming 2018). Given S1’s role in encoding somatosensory afferent input from button press movements, it may contribute primarily to confirmatory post-movement feedback about the successful execution of each chosen motor-report movement, independent of the correctness of the preceding sensorimotor decision or the strength of evidence on which it was based.

### General Conclusions

Our results align with models proposing that metacognitive performance monitoring networks, including the cingulate cortex, are more engaged under decision uncertainty, and that outcome feedback is evaluated relative to internal expectations, rather than in isolation (Kepecs et al. 2008; Kiani et al. 2014; Vassena et al. 2020). The present results show that these signals are also expressed in premotor cortical ROIs outside of the cingulate cortex (An et al. 2019; Ramakrishnan et al. 2017; FitzGerald et al. 2013; Bang and Fleming 2018; Ramkumar et al. 2016; So and Stuphorn 2012; 2016; Levy et al. 2020). This also aligns with models of embodied decision-making and of metacognitive evaluation and hierarchical Bayesian inference, where cortical activity reflects adjustments to internal models based on outcome confirmation or violation (Friston et al. 2013; O’Reilly 2013).

These findings support the view that the brain continuously monitors and evaluates its own decisions beyond a simple binary correct/incorrect categorization. Instead, the evidence-dependent +RPE-like modulations of beta and alpha band responses to identical correct Feedback-Beep signals, combined with the participants’ reports of evidence-dependent changes in their level of confidence about their decisions, indicated that the dynamics of outcome feedback processing was strongly influenced by internal inferential processes, such as decision confidence and expectations for decision outcomes. The +RPE-like beta- and alpha-band post-Feedback-Beep signals were widely broadcast across many ROIs in these sensorimotor decision-making tasks, where they could be used to update the dynamical state of neural circuits implicated in many aspects of the sensorimotor decision-making process, including belief in and accumulation of color sensory evidence, decision thresholds, decision confidence and action-selection processes (Dehaene et al. 2021; Drugowitsch et al. 2014; Pouget et al. 2016; Donner et al. 2007; 2009; Lak et al. 2017; Tsunada et al. 2019).

It is also noteworthy that these +RPE-like signals were generated at the end of the correct trials even though the participants received no overt rewards for successful performance. There was no monetary incentive or accumulating point score to motivate the participants, and they did not receive anything resembling the hedonic or appetitive physical rewards (e.g., juice drops, food pellets) typically delivered to non-human participants in such tasks, against which the participants could compare the experienced outcome to calculate a concrete “reward prediction error”. Instead, the rewarding value of correct decisions in the present tasks may have resulted from some innate value that the participants attributed to the successful completion of a trial, especially when it was difficult, or even from more pragmatic “rewards” such as the knowledge that they were one trial closer to completing a trial block.

This reinforces the proposal that these +RPE-like responses reflected computations based on internal metacognitive processes informed by the participants’ assessment of the likely correctness of their decisions as a function of different levels of sensory evidence and associated expectations of the resulting likely outcomes. Moreover, they were expressed while the participants performed deterministic tasks in which the decision outcomes was based entirely on their evaluation of the sensory evidence. Their confidence about their decisions was based entirely on their prior experience in a task environment with veridical outcome feedback, rather than in an environment in which experienced outcomes and uncertainty about outcomes were imposed externally by probabilistic reward schedules.

## Acknowledgements

This research was supported by grants from the Canadian Institutes of Health Research (CIHR MOP-97944 [J.K.] and MOP-142220 [J.K., S.B.]); a doctoral studentship from the NSERC-CREATE: Complex Dynamics of Brain and Behaviour program; a scholarship from the McGill University Integrated Program in Neuroscience; and a doctoral studentship from the Fonds de Recherche du Québec (FRQ 318042, DOI: https://doi.org/10.69777/318042).

## Supplementary Figures

**Supplemental Figure 1:**
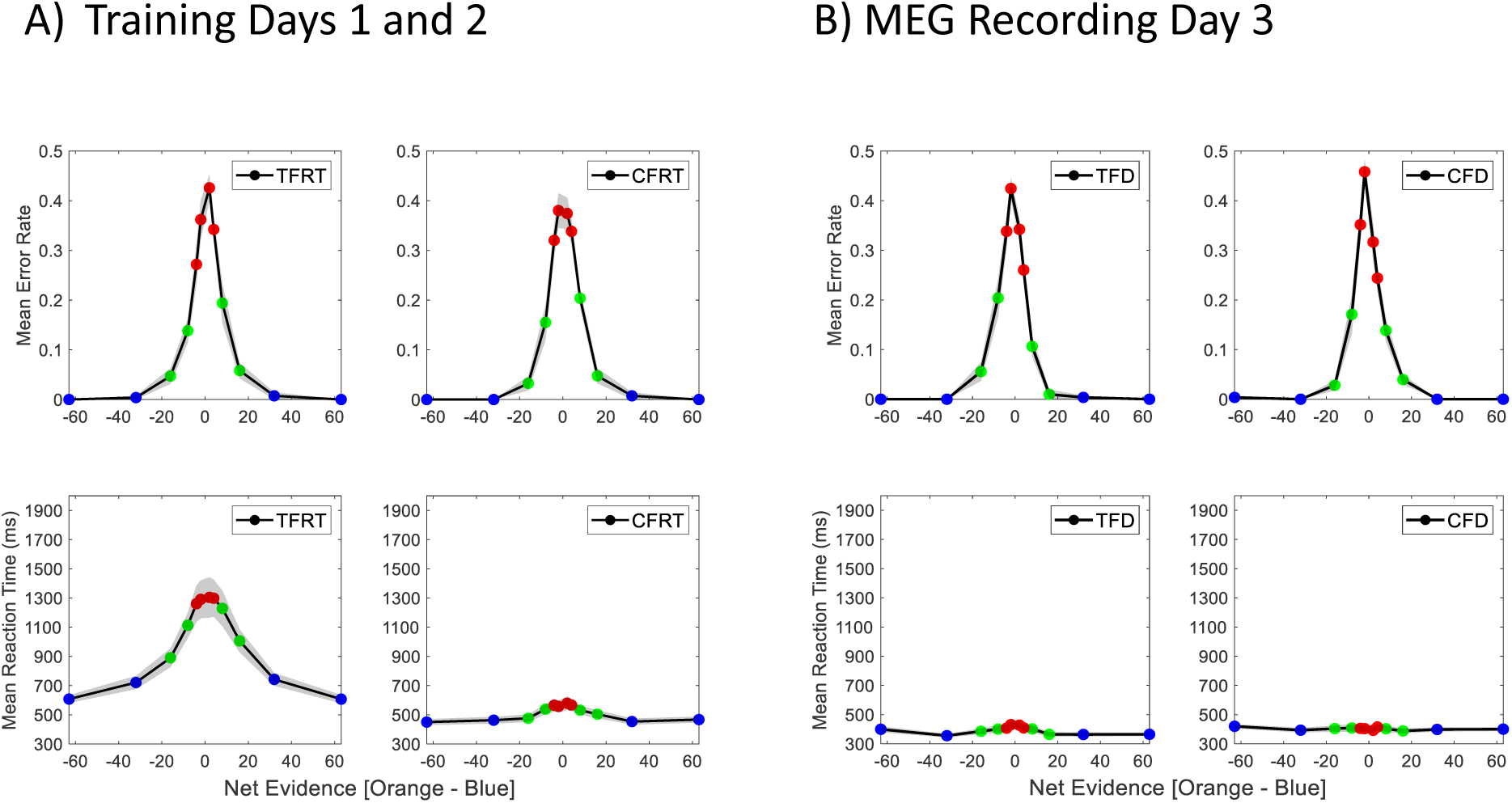
Grand mean error rates (±s.e.m.) (top row) and grand mean reaction times (±s.e.m.) (bottom row) in correct trials with different levels of color evidence (Strong: blue dots; Intermediate: green dots; Weak: red dots), while performing the TFRT and CFRT tasks during the two training sessions (A), and the TFD and CFD tasks during the MEG recording session (B). Across all 4 tasks, error rates increased systematically as evidence strength decreased from Strong to Intermediate and Weak. Decision-report buttonpress RTs increased systematically when CKBs with weaker color evidence appeared at the start of the TargetCKBRT epoch in the TFRT task, indicating that the participants took longer to choose the dominant color of the CKBs as net color evidence in the CKBs decreased. This effect of color evidence on RTs largely disappeared in the CFRT task, when the CKBs appeared for 2000-3000ms before the colored Target Cues. In contrast, RTs were nearly identical and unmodulated by color evidence strength in the CFD and TFC tasks during MEG recording day 3, because of the second 2000-3000ms delay imposed in those tasks before the appearance of the Go/RT cues.

**Supplemental Figure 2.**
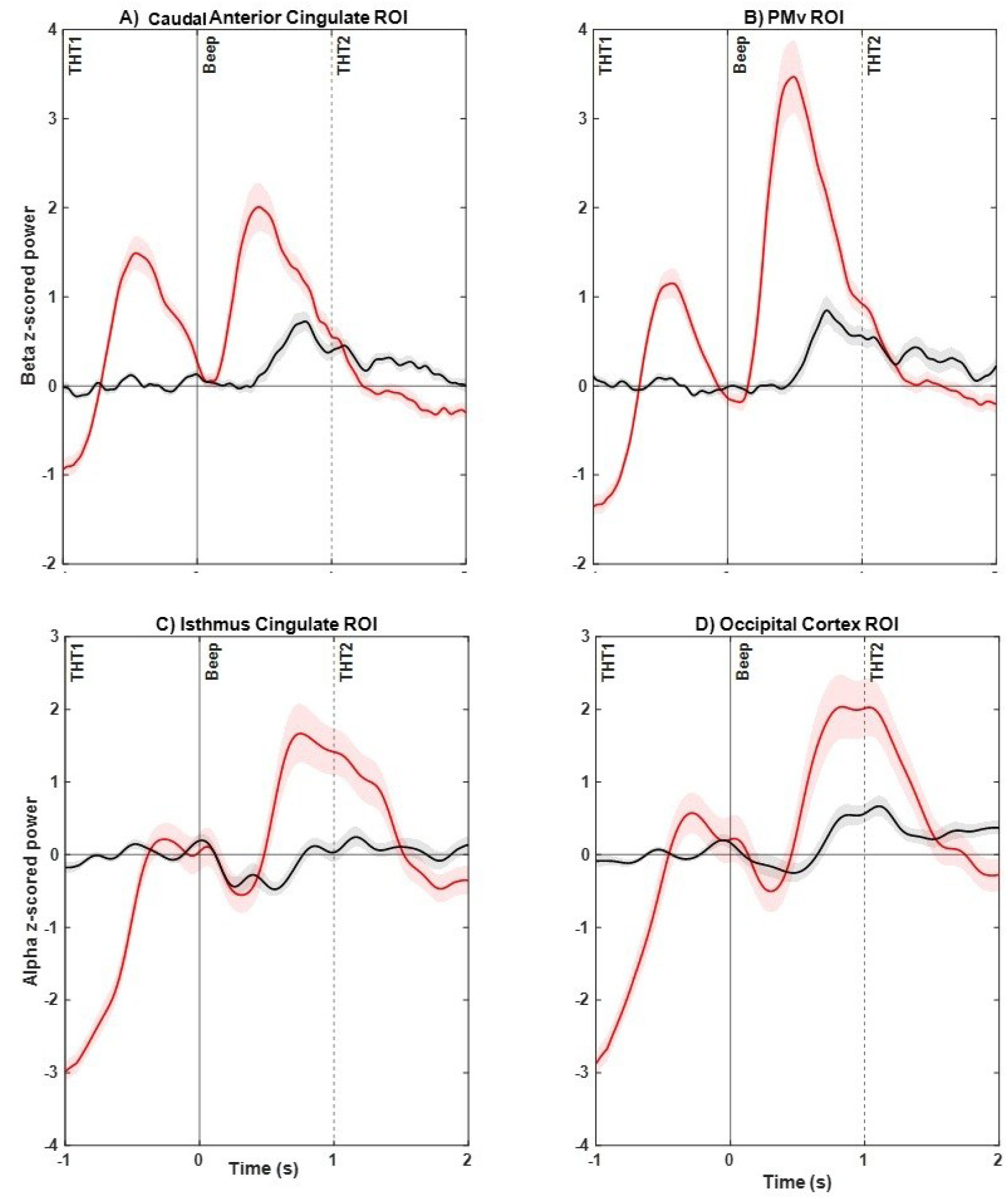
A) Grand mean (± s.e.m.) beta-band response profiles in “correct” Random-Beep trials (black) and in correct TFD/CFD Weak-evidence trials (red) in the Caudal Anterior Cingulate ROI, aligned to the onset of the Beep tones (time 0sec). **B)** Grand mean (± s.e.m.) beta-band response profiles in the PMv ROI. **C)** Grand mean (± s.e.m.) alpha-band response profiles in the Isthmus Cingulate ROI. **C)** Grand mean (± s.e.m.) alpha-band response profiles in the Occipital ROI. **B, C, D)** Same format as in **(A)**.

**Supplemental Figure 3.**
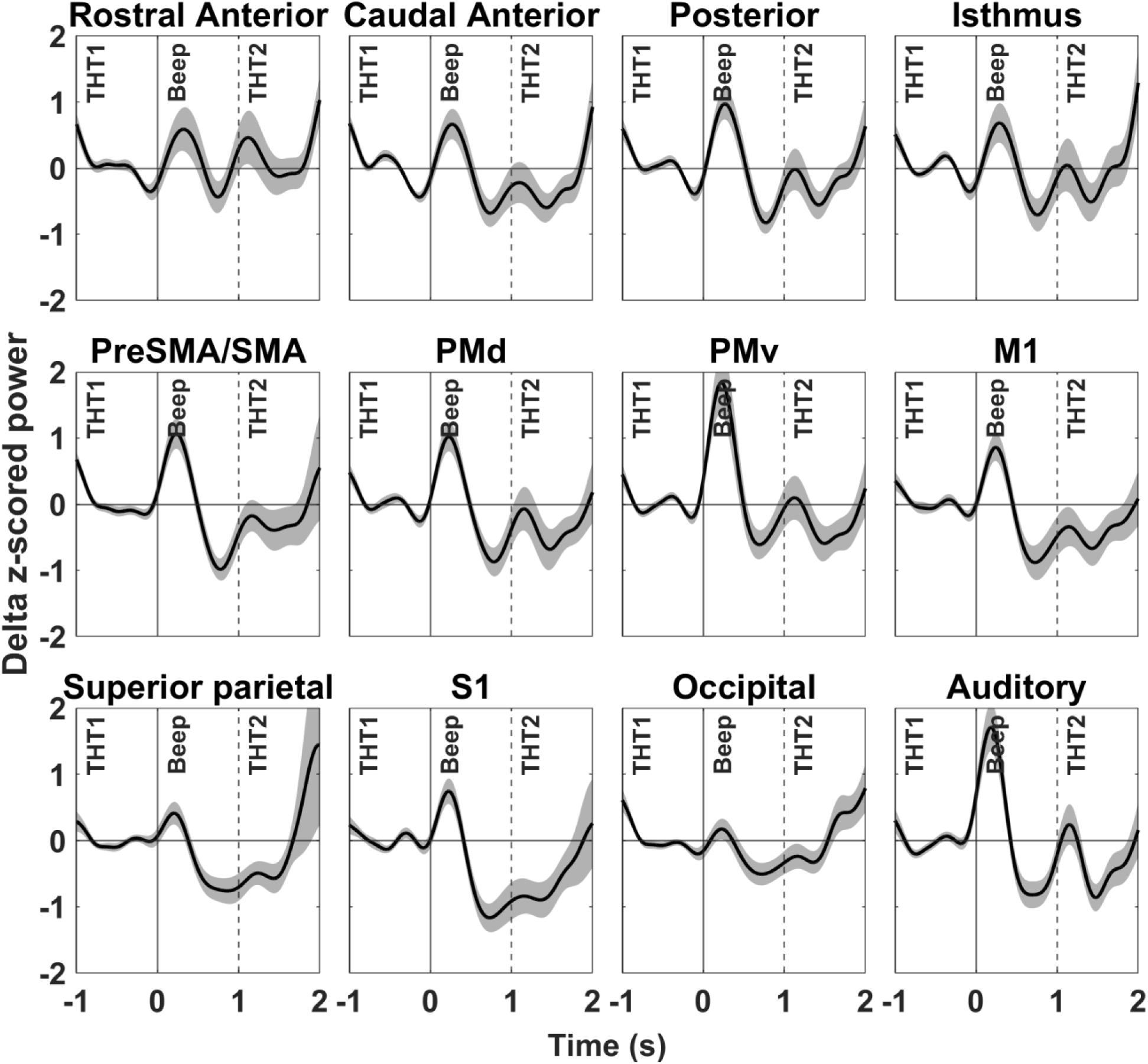
Grand-mean z-score-normalized (±s.e.m.) delta (2-4 Hz) power in four cingulate ROIs (top row) and 8 other cortical ROIs (middle and bottom rows) in response to the “correct” 1500 Hz Random-Beep auditory tones. Same format as Figure 6.

**Supplemental Figure 4.**
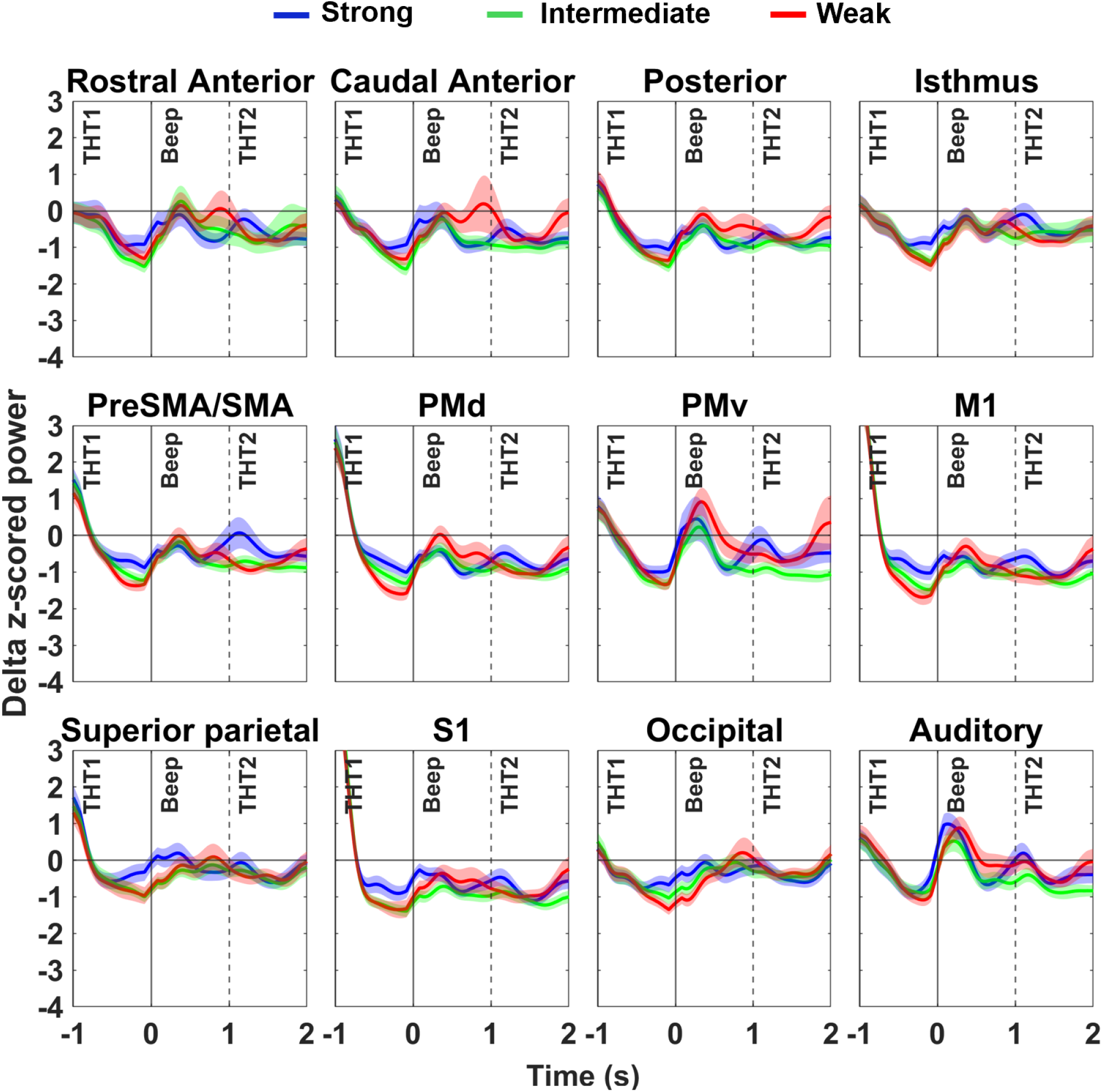
Grand-mean z-score-normalized (±s.e.m.) delta (2-4 Hz) power in four cingulate ROIs (top row) and 8 other cortical ROIs (middle and bottom rows) across the three post-decision epochs (THT1, Beep, THT2), aligned to Feedback-Beep onset (time 0). Only trials with correct decisions were included. Same format as Figure 8.

## Supplemental Tables

**Supplemental Table 1.** Summary of the design of the four tasks, and the neural computations expected during each epoch of trial before the selected target is reported by a mouse button press.

| Task | Cue Order | Expected decision processes |  | When to report final decision |
| --- | --- | --- | --- | --- |
|  |  | First Cue | Second Cue |  |
| TFRT | Targets → CKB | Identify target color-location conjunctions | Identify dominant color of CKB and select target with the same color | Report as soon as the target decision is made after the CKB appears |
| CFRT | CKB → Targets | Identify dominant color of CKB | Identify target color-location conjunctions and select target with the same color as the CKB | Report as soon as the target decision is made after the colored targets appear |
| TFD | Targets → CKB | Identify target color-location conjunctions | Identify dominant color of CKB and select target with the same color | Delay target decision report until an explicit Go Cue appears |
| CFD | CKB → Targets | Identify dominant color of CKB | Identify target color-location conjunctions and select target with the same color as the CKB | Delay target decision report until an explicit Go Cue appears |

**Supplemental Table 2.** Grand mean (± s.e.m.) z-scored magnitude of the beta-band response to correct Feedback-Beep cues in trials with different evidence strength in defined ROIs, and in response to “correct” Random-Beep tones.

| Evidence Strength: | Strong | Intermediate | Weak | Random-Beep |
| --- | --- | --- | --- | --- |
| Cingulate ROI: |  |  |  |  |
| RAC | $0.98 \pm 0.04$ | $2.01 \pm 0.16$ | $2.47 \pm 0.19$ | $1.09 \pm 0.08$ |
| CAC | $1.91 \pm 0.07$ | $2.34 \pm 0.21$ | $2.96 \pm 0.25$ | $1.36 \pm 0.13$ |
| PC | $1.14 \pm 0.07$ | $1.65 \pm 0.11$ | $2.03 \pm 0.18$ | $1.73 \pm 0.16$ |
| IC | $1.10 \pm 0.06$ | $1.42 \pm 0.10$ | $1.83 \pm 0.10$ | $1.38 \pm 0.15$ |
| Cortical ROI: |  |  |  |  |
| PreSMA/SMA | $1.15 \pm 0.09$ | $1.73 \pm 0.16$ | $2.19 \pm 0.25$ | $1.54 \pm 0.22$ |
| PMd | $1.33 \pm 0.10$ | $1.93 \pm 0.15$ | $2.41 \pm 0.23$ | $1.80 \pm 0.26$ |
| PMv | $1.56 \pm 0.12$ | $3.46 \pm 0.30$ | $4.86 \pm 0.39$ | $1.60 \pm 0.18$ |
| M1 | $1.22 \pm 0.10$ | $1.66 \pm 0.15$ | $1.78 \pm 0.17$ | $1.99 \pm 0.26$ |
| SPC | $0.93 \pm 0.08$ | $1.43 \pm 0.10$ | $1.81 \pm 0.13$ | $1.44 \pm 0.18$ |
| S1 | $1.28 \pm 0.11$ | $1.66 \pm 0.13$ | $1.93 \pm 0.19$ | $1.89 \pm 0.26$ |
| OCC | $0.94 \pm 0.09$ | $1.43 \pm 0.10$ | $2.25 \pm 0.18$ | $1.14 \pm 0.13$ |
| AC | $1.39 \pm 0.10$ | $2.20 \pm 0.14$ | $2.67 \pm 0.13$ | $1.52 \pm 0.25$ |

**Abbreviations:**
ROI: region of interest; RAC: rostral anterior cingulate; CAC: caudal anterior cingulate; PC: posterior cingulate; IC: isthmus cingulate; preSMA/SMA: combined pre-supplemental motor area and supplemental motor area; PMd: dorsal premotor cortex; PMv: ventral premotor cortex; M1: primary motor cortex; SPC: superior parietal cortex; S1: primary somatosensory cortex; OCC: occipital cortex; AC: auditory cortex.

**Supplemental Table 3.** Grand mean (± s.e.m.) z-scored magnitude of the alpha-band suppression response to correct Feedback-Beep cues in trials with different evidence strength in defined ROIs, and in response to “correct” Random-Beep tones. Same format as Supplemental Table 2.

| Evidence Strength: | Strong | Intermediate | Weak | Random-Beep |
| --- | --- | --- | --- | --- |
| Cingulate ROI: |  |  |  |  |
| RAC | -1.34 ± 0.12 | -1.28 ± 0.10 | -1.02 ± 0.09 | -1.55 ± 0.12 |
| CAC | -1.53 ± 0.11 | -1.37 ± 0.12 | -1.58 ± 0.12 | -1.39 ± 0.12 |
| PC | -1.33 ± 0.12 | -1.48 ± 0.10 | -1.39 ± 0.12 | -1.54 ± 0.12 |
| IC | -1.78 ± 0.16 | -1.82 ± 0.17 | -1.27 ± 0.13 | -1.44 ± 0.12 |
| Cortical ROI: |  |  |  |  |
| PreSMA/SMA | -1.44 ± 0.10 | -1.64 ± 0.14 | -1.92 ± 0.16 | -1.46 ± 0.16 |
| PMd | -1.66 ± 0.13 | -1.84 ± 0.16 | -2.61 ± 0.33 | -1.45 ± 0.15 |
| PMv | -1.90 ± 0.15 | -1.64 ± 0.13 | -2.06 ± 0.17 | -2.12 ± 0.20 |
| M1 | -1.69 ± 0.16 | -1.91 ± 0.18 | -2.78 ± 0.31 | -1.76 ± 0.16 |
| SPC | -1.70 ± 0.17 | -1.88 ± 0.20 | -1.47 ± 0.16 | -1.37 ± 0.15 |
| S1 | -1.93 ± 0.21 | -1.95 ± 0.22 | -2.51 ± 0.27 | -1.93 ± 0.20 |
| OCC | -1.70 ± 0.17 | -1.65 ± 0.16 | -1.47 ± 0.16 | -1.13 ± 0.11 |
| AC | -3.10 ± 0.26 | -2.98 ± 0.22 | -2.82 ± 0.21 | -3.35 ± 0.26 |

**Supplemental Table 4.** Grand mean (± s.e.m.) z-scored magnitude of the alpha-band rebound response to correct Feedback-Beep cues in trials with different evidence strength in defined ROIs, and in response to “correct” Random-Beep tones. Same format as Supplemental Table 2.

| Evidence Strength: | Strong | Intermediate | Weak | Random-Beep |
| --- | --- | --- | --- | --- |
| Cingulate ROI: |  |  |  |  |
| RAC | $1.39 \pm 0.11$ | $2.06 \pm 0.16$ | $2.15 \pm 0.14$ | $2.27 \pm 0.17$ |
| CAC | $1.24 \pm 0.10$ | $2.14 \pm 0.12$ | $2.04 \pm 0.18$ | $1.78 \pm 0.14$ |
| PC | $1.40 \pm 0.11$ | $2.18 \pm 0.20$ | $2.21 \pm 0.18$ | $1.96 \pm 0.20$ |
| IC | $1.56 \pm 0.17$ | $2.66 \pm 0.28$ | $3.34 \pm 0.23$ | $1.99 \pm 0.13$ |
| Cortical ROI: |  |  |  |  |
| PreSMA/SMA | $2.08 \pm 0.39$ | $1.61 \pm 0.14$ | $1.74 \pm 0.20$ | $1.97 \pm 0.20$ |
| PMd | $1.94 \pm 0.19$ | $1.91 \pm 0.15$ | $2.10 \pm 0.21$ | $1.97 \pm 0.21$ |
| PMv | $1.88 \pm 0.16$ | $2.41 \pm 0.22$ | $2.37 \pm 0.26$ | $2.19 \pm 0.29$ |
| M1 | $1.94 \pm 0.23$ | $1.90 \pm 0.16$ | $2.06 \pm 0.20$ | $2.47 \pm 0.27$ |
| SPC | $1.97 \pm 0.21$ | $2.43 \pm 0.27$ | $2.82 \pm 0.26$ | $2.10 \pm 0.24$ |
| S1 | $2.12 \pm 0.28$ | $2.26 \pm 0.21$ | $2.41 \pm 0.27$ | $2.50 \pm 0.31$ |
| OCC | $1.57 \pm 0.19$ | $3.17 \pm 0.36$ | $3.76 \pm 0.30$ | $1.88 \pm 0.18$ |
| AC | $2.01 \pm 0.18$ | $3.02 \pm 0.26$ | $3.23 \pm 0.27$ | $2.22 \pm 0.28$ |

**Supplemental Table 5.** Grand mean (± s.e.m.) z-scored magnitude of the post-report-movement beta-band rebound response during the THT1 epoch in trials with different evidence strength in defined ROIs. Similar format as Supplemental Table 2.

| Evidence Strength: | Strong | Intermediate | Weak |
| --- | --- | --- | --- |
| RAC | $0.33 \pm 0.06$ | $0.52 \pm 0.06$ | $0.27 \pm 0.08$ |
| CAC | $0.97 \pm 0.14$ | $1.19 \pm 0.17$ | $0.83 \pm 0.14$ |
| PC | $1.56 \pm 0.26$ | $1.42 \pm 0.20$ | $1.16 \pm 0.18$ |
| IC | $0.75 \pm 0.11$ | $0.75 \pm 0.11$ | $0.67 \pm 0.19$ |
| PreSMA/SMA | $2.10 \pm 0.32$ | $1.94 \pm 0.28$ | $1.54 \pm 0.24$ |
| PMd | $2.39 \pm 0.38$ | $2.27 \pm 0.32$ | $1.73 \pm 0.27$ |
| PMv | $1.06 \pm 0.15$ | $0.87 \pm 0.15$ | $0.52 \pm 0.13$ |
| M1 | $2.69 \pm 0.49$ | $2.45 \pm 0.37$ | $1.93 \pm 0.30$ |
| SPC | $0.90 \pm 0.27$ | $1.07 \pm 0.22$ | $0.70 \pm 0.19$ |
| S1 | $1.98 \pm 0.42$ | $1.96 \pm 0.34$ | $1.47 \pm 0.28$ |
| OCC | $1.49 \pm 0.18$ | $1.79 \pm 0.18$ | $1.66 \pm 0.21$ |
| AC | $0.62 \pm 0.09$ | $0.53 \pm 0.09$ | $0.27 \pm 0.10$ |

**Supplemental Table 6.** Grand mean (± s.e.m.) z-scored magnitude of the post-report-movement alpha-band rebound response during the THT1 epoch in trials with different evidence strength in defined ROIs. Same format as Supplemental Table 5.

| Evidence Strength: | Strong | Intermediate | Weak |
| --- | --- | --- | --- |
| RAC | $0.77 \pm 0.36$ | $0.59 \pm 0.39$ | $-0.16 \pm 0.36$ |
| CAC | $0.91 \pm 0.34$ | $1.01 \pm 0.47$ | $0.08 \pm 0.38$ |
| PC | $1.49 \pm 0.50$ | $1.16 \pm 0.44$ | $-0.37 \pm 0.50$ |
| IC | $1.55 \pm 0.56$ | $1.95 \pm 0.61$ | $-0.58 \pm 0.55$ |
| PreSMA/SMA | $1.69 \pm 0.38$ | $1.42 \pm 0.39$ | $0.96 \pm 0.41$ |
| PMd | $1.76 \pm 0.48$ | $1.36 \pm 0.43$ | $1.20 \pm 0.55$ |
| PMv | $1.59 \pm 0.35$ | $1.28 \pm 0.42$ | $0.51 \pm 0.37$ |
| M1 | $1.67 \pm 0.53$ | $1.07 \pm 0.49$ | $0.98 \pm 0.59$ |
| SPC | $1.24 \pm 0.52$ | $1.53 \pm 0.65$ | $-0.80 \pm 0.52$ |
| S1 | $1.83 \pm 0.55$ | $1.28 \pm 0.53$ | $0.55 \pm 0.52$ |
| OCC | $2.13 \pm 0.71$ | $1.95 \pm 0.61$ | $-0.34 \pm 0.59$ |
| AC | $1.58 \pm 0.46$ | $1.16 \pm 0.50$ | $-0.02 \pm 0.53$ |

**Supplemental Table 7.** Mean difference in z-scored post-Feedback-Beep response magnitude across all ROIs (and two-tailed one-sample t-test results) in correct trials with different levels of color evidence strength in defined ROIs (Supplemental Tables 2, 3, 4).

| Comparison: | Different levels of evidence strength |  |  |
| --- | --- | --- | --- |
| Evidence: | W-S | I-S | W-I |
| Response Component: |  |  |  |
| Beta rebound | 1.26<br>(**<0.001**) | 0.77<br>(**<0.001**) | 0.48<br>(**<0.001**) |
| Alpha suppression | -0.14<br>(0.38) | -0.02<br>(0.71) | -0.12<br>(0.39) |
| Alpha rebound | 0.76<br>(**0.003**) | 0.56<br>(**0.007**) | 0.21<br>(0.011) |

p-value format: **p<0.01**; 0.01<p<0.05; p>0.05.

**Supplemental Table 8.** Mean difference in z-scored pre-Feedback-Beep THT1 rebound magnitude across all 12 ROIs (and two-tailed one-sample t-test results) in correct trials with different levels of color evidence strength (Supplemental Tables 5, 6).

| Comparison: | Different levels of evidence strength |  |  |
| --- | --- | --- | --- |
| Evidence: | W-S | I-S | W-I |
| Response component: |  |  |  |
| Beta THT1 rebound | -0.34<br>(**0.002**) | -0.01<br>(0.90) | -0.33<br>(**<0.001**) |
| Alpha THT1 rebound | -1.3<br>(**<0.001**) | -0.20<br>(0.05) | -1.14<br>(**<0.001**) |

p-value format: **p<0.01**; 0.01<p<0.05; p>0.05.

**Supplemental Table 9.** Grand mean magnitude (in arbitrary z-score units) of post-Feedback-Beep beta and alpha rebound responses in correct trials across all 12 ROIs in three consecutive trial sub-blocks within the 4 trial blocks in the TFD and CFD tasks (and two-tailed one-sample t-test results comparing the rebound magnitudes in SB2 and SB3 against the values in SB1).

| Sub-Blocks | SB1 | SB2 | SB3 |
| --- | --- | --- | --- |
| Beta Rebound |  |  |  |
| Strong | 2.144 | 1.922<br>(0.013) | 2.254<br>(0.159) |
| Intermediate | 2.863 | 2.838<br>(0.799) | 2.876<br>(0.921) |
| Weak | 3.299 | 3.330<br>(0.764) | 3.494<br>(0.168) |
| Alpha Rebound |  |  |  |
| Strong | 3.247 | 2.875<br>(0.009) | 3.757<br>(**<0.001**) |
| Intermediate | 3.570 | 3.817<br>(0.123) | 3.577<br>(0.968) |
| Weak | 3.626 | 4.110<br>(0.014) | 3.682<br>(0.821) |

p-value format: **p<0.01**; 0.01<p<0.05; p>0.05.

**Supplemental Table 10.** Grand mean (± s.e.m.) z-scored magnitude of the delta-band rebound response to correct Feedback-Beep cues in trials with different evidence strength in defined ROIs, and in response to “correct” Random-Beep tones. Same format as Supplemental Table 2.

| Evidence Strength: | Strong | Intermediate | Weak | Random-Beep |
| --- | --- | --- | --- | --- |
| Cingulate ROI: |  |  |  |  |
| RAC | 1.74 ± 0.29 | 1.46 ± 0.32 | 2.02 ± 0.50 | 1.94 ± 0.42 |
| CAC | 1.27 ± 0.19 | 1.38 ± 0.15 | 2.50 ± 0.80 | 1.09 ± 0.25 |
| PC | 1.11 ± 0.18 | 1.36 ± 0.12 | 1.92 ± 0.33 | 1.27 ± 0.29 |
| IC | 1.69 ± 0.34 | 1.45 ± 0.18 | 1.76 ± 0.32 | 1.56 ± 0.40 |
| Cortical ROI: |  |  |  |  |
| PreSMA/SMA | 1.75 ± 0.40 | 1.31 ± 0.23 | 1.76 ± 0.29 | 0.76 ± 0.25 |
| PMd | 1.27 ± 0.23 | 1.30 ± 0.14 | 1.98 ± 0.27 | 1.08 ± 0.37 |
| PMv | 1.80 ± 0.49 | 1.09 ± 0.11 | 2.05 ± 0.39 | 1.12 ± 0.35 |
| M1 | 1.17 ± 0.23 | 1.20 ± 0.12 | 1.69 ± 0.19 | 0.79 ± 0.32 |
| SPC | 1.35 ± 0.27 | 1.66 ± 0.20 | 1.96 ± 0.25 | 0.52 ± 0.20 |
| S1 | 1.19 ± 0.26 | 1.37 ± 0.14 | 1.81 ± 0.25 | 0.21 ± 0.27 |
| OCC | 1.58 ± 0.22 | 1.66 ± 0.19 | 2.34 ± 0.42 | 0.90 ± 0.18 |
| AC | 1.92 ± 0.37 | 1.28 ± 0.13 | 2.16 ± 0.29 | 0.96 ± 0.30 |

